# SIM2s coordinates programmed mitophagy with respiratory chain supercomplex remodeling during mammary epithelial differentiation

**DOI:** 10.64898/2026.09.14.750442

**Authors:** Ramsey M. Jenschke, Madelyn Spooner-Harris, Steven W. Wall, Lilia Sanchez, Jessica Epps, Sophia Schutmaat, Kaitlin P. Kelly, Tyler Frey, Monique Rijnkels, Weston W. Porter

## Abstract

Lactation requires mammary epithelial cells (MECs) to rapidly expand mitochondrial function while remodeling the mitochondrial population that supports milk synthesis and secretion. Programmed mitophagy is required for MEC differentiation, yet why mitochondrial turnover is necessary during this developmental transition remains poorly understood. Using Mito-QC reporter mice, we identified developmentally regulated changes in mitolysosome burden across the transition from late pregnancy to lactation that were altered by mammary-specific gain or loss of the bhlh/PAS protein, SIM2s (single-minded 2 s). In differentiating HC11 cells, mitochondrial turnover was accompanied by increased assembly and activity of respiratory supercomplexes containing complexes I, III, and IV. Depletion of PRKN prevented acquisition of this differentiation-associated respiratory profile and impaired lactogenic differentiation. SIM2s co-migrated with higher-order respiratory assemblies, and loss of SIM2s reduced supercomplex assembly and activity in HC11 cells and mammary tissue. SIM2s also localized in close proximity to complex III in differentiated mammary epithelium, whereas loss of SIM2s reduced proximity between complexes III and IV. Together, these findings support a model in which SIM2s coordinates PRKN-dependent mitochondrial turnover with respiratory-chain remodeling during MEC differentiation. Our results suggest that programmed mitophagy does more than remove mitochondria during development; it contributes to establishment of a mitochondrial population with a respiratory-chain architecture suited to the emerging differentiated state.

## Introduction

The transition from pregnancy to lactation requires profound structural and metabolic adaptation in the mammary gland [1,2]. At parturition, mammary epithelial cells (MECs) rapidly acquire the capacity to continuously synthesize and secrete large quantities of proteins, lipids, and carbohydrates. Mitochondrial content and oxidative capacity increase as lactation begins, yet lactogenic differentiation also engages programmed mitochondrial turnover. The purpose of this turnover in a cell state that requires greater mitochondrial output remains poorly defined. Specifically, it is not known whether mitophagy simply preserves mitochondrial quality or actively contributes to establishing a mitochondrial population with functions required by the differentiated cell [3–7]. Indeed, mitochondrial dysfunction has recently been linked to delays in milk secretion in insulin resistant women, suggesting the importance of a mitochondrial population capable of the adaption required for lactation [8,9].

Mitophagy is the selective autophagic degradation of mitochondria and is commonly studied after mitochondrial damage or depolarization. Programmed mitophagy is instead engaged by developmental signals as cells acquire a new identity. In cardiomyocytes and myoblasts, mitochondrial turnover supports the transition to an oxidative state, whereas mitophagy promotes a glycolytic transition during retinal ganglion cell differentiation [10–12]. These different outcomes suggest that programmed mitophagy does not impose a common metabolic endpoint. Rather, it may remove mitochondria that are no longer suited to the emerging cell state and permit their replacement with organelles adapted to the new metabolic program. Prior studies have largely defined changes in mitochondrial abundance, morphology, biogenesis, or respiration, but the structural features that distinguish the replacement mitochondrial population remain unclear [12–14]. Thus, the functional reason that mitochondrial turnover is required for differentiation has not been resolved. More broadly, selective autophagy may serve as an active mechanism for rebuilding organelle systems during differentiation rather than simply removing damaged material.

We previously identified SIM2s, the short isoform of the bHLH/PAS factor single-minded 2, as a breast tumor suppressor and important regulator of mammary epithelial differentiation and lactation [15–26]. We found that loss of *Sim2* impairs mammary gland development and lactation, whereas moderate overexpression of SIM2s promotes differentiation and prolongs MEC function [17–19,27]. Although SIM2s is classically defined as a transcription factor, we subsequently demonstrated that it also localizes to mitochondria, where it promotes PRKN (parkin RBR E3 ubiquitin protein ligase) recruitment and mitochondrial turnover during MEC differentiation [28,29]. Disruption of autophagy or depletion of PRKN impairs mitochondrial respiration, cell survival, and differentiation-dependent gene expression [17–19,28]. Together, these findings position SIM2s as a potential link between developmental signaling and the mitochondrial remodeling required for lactation. However, they did not define the functional mitochondrial state produced by SIM2s- and PRKN-dependent turnover.

A potential functional output of programmed mitophagy is reorganization of the mitochondrial respiratory chain (MRC). Complexes I, III, and IV can assemble into higher-order respiratory supercomplexes (SCs), whose abundance and composition vary with metabolic state. Although the functional consequences of SC assembly remain debated, these structures can stabilize individual complexes, support respiratory-chain maturation, and influence electron transfer and redox output [30,31]. Increased SC abundance has been observed during adipogenic differentiation, mitochondrial protein assemblies remodel during neurogenesis, and the PINK1-PRKN pathway can selectively regulate turnover of respiratory-chain proteins [32–34]. However, to our knowledge, no study has directly shown that programmed mitophagy is required to establish a differentiation-associated MRC supercomplex organization [35,36]. In breast cancer cells, mitochondrial SIM2s associates with MRC proteins, supports CIII stability and SC assembly, and promotes oxidative phosphorylation [29,33]. Thus, SIM2s provides a potential mechanistic link between mitochondrial turnover and respiratory-chain maturation during normal differentiation.

Based on these findings, we investigated whether programmed mitophagy produces a mitochondrial population with a distinct MRC organization during the transition to lactation and whether SIM2s coordinates these processes. *Mito-QC* reporter mice [37–39] identified developmentally patterned changes in mitolysosome burden that were altered by mammary-specific gain or loss of *Sim2*. In HC11 cells and mammary tissue, differentiation was accompanied by increased assembly and activity of CI-, CIII-, and CIV-containing SCs. Depletion of PRKN prevented acquisition of this respiratory profile and impaired lactogenic differentiation, whereas loss of SIM2s reduced SC assembly and activity. SIM2s also co-migrated with higher-order respiratory assemblies and localized near CIII in differentiated mammary epithelium. Together, these findings identify respiratory-chain remodeling as a functional output of programmed mitophagy. They support a model in which mitochondrial turnover permits acquisition of a respiratory-chain organization suited to the emerging differentiated state, with SIM2s coordinating mitochondrial renewal and respiratory maturation.

## Results

### Programmed mitophagy increases during the transition into lactation

Our previous HC11 studies established programmed mitophagy as a requirement for lactogenic differentiation. We first defined the developmental timing of mitochondrial turnover in vivo using the Rosa26-mCherry-GFP-FIS1^101-152 (*Mito-QC*) reporter mouse [37,40]. In this model, mitochondria display both GFP and mCherry fluorescence under basal conditions, whereas delivery to the acidic lysosomal compartment quenches GFP fluorescence while preserving mCherry, generating mCherry-positive/GFP-negative puncta that mark mitolysosomes. Mammary glands were collected from virgin mice and across pregnancy, lactation, and involution to determine how mitochondrial turnover changes during functional differentiation of the gland (Figure 1A-B).

**Figure 1.**
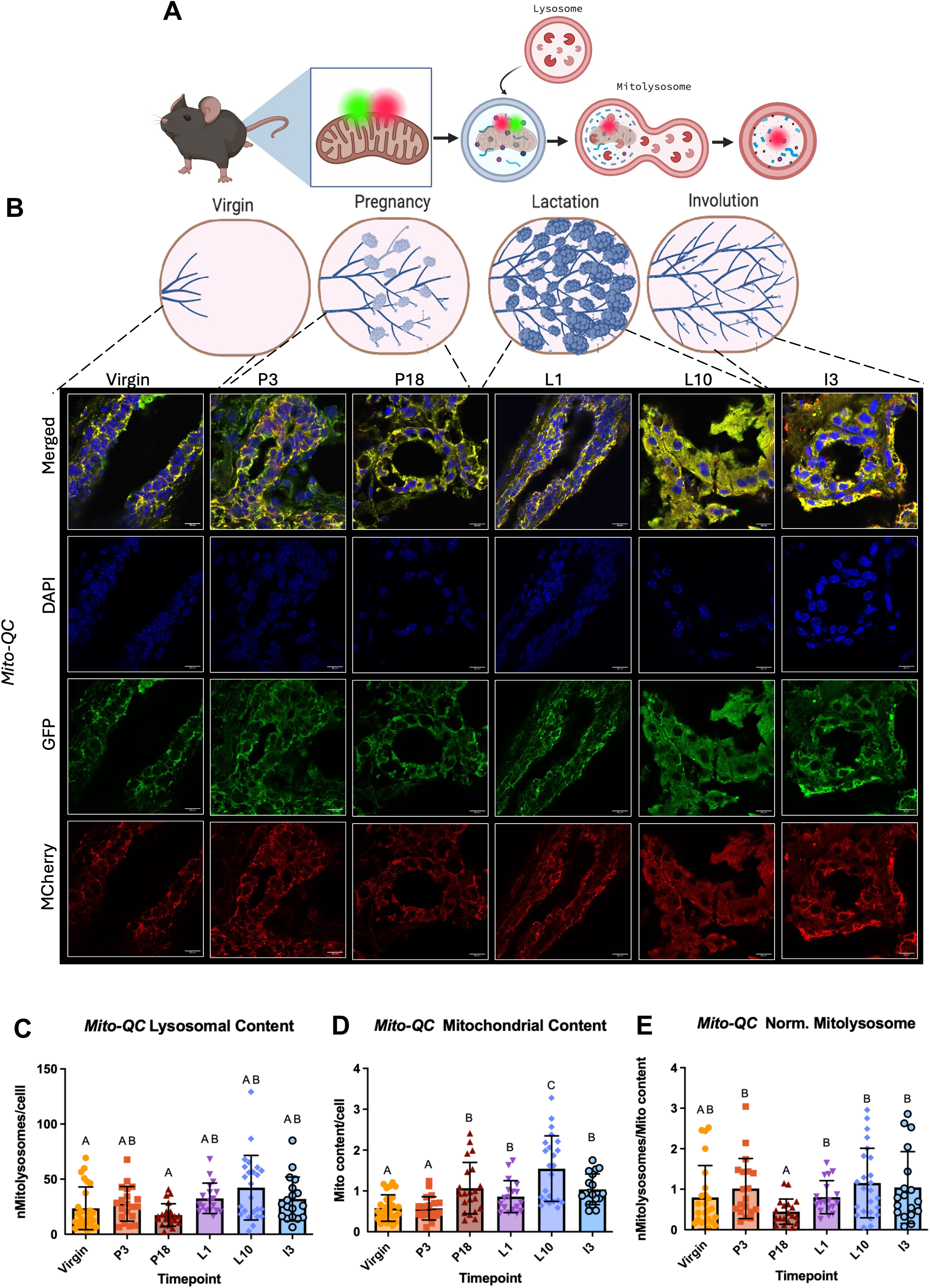
*Mito-QC* mouse model mammary development demonstrates temporally modulated mitolysosome formation (**A**) *Mito*-QC reporter mice were sacrificed at key developmental timepoints, and mammary glands were processed for confocal analysis. Created with BioRender.com. (**B**) Representative images of *Mito-QC* mammary glands taken at virgin (in estrus), pregnancy days 3 (P3) and 18 (P18), lactation days 1 (L1) and 10 (L10), and involution day 3 (I3) and stained for immunofluorescence with GFP and mCherry antibodies. Images were taken using 100x objective split into merged (GFP+, mCherry+, and DAPI) and individual channels. Scale bars 10uM. All timepoints have *n* = 4–6 mice with 3-6 images per mouse. (**C**) Quantification of mitolysosome number (mCherry^+^GFP^−^ puncta) normalized for cell count per image. (**D**) Quantification of mitochondrial content (mean GFP intensity) normalized for cell count. (**E**) Quantification of mitolysosomes controlled for mitochondrial content, normalized to cell count. All data are expressed as the mean ± SD. Dots represent individual images. For all variables with the same letter, the difference between the means is not statistically significant. Significance was calculated using multiple two-tailed unpaired or Welch’s *t*-tests. *P* < 0.05 was considered statistically significant.

To distinguish changes in mitochondrial degradation from changes in the size of the mitochondrial population, we quantified mitolysosome burden and mitochondrial content separately. Mitolysosome burden was determined from mCherry-positive puncta, whereas mean GFP fluorescence was used as a measure of mitochondrial content, with both measurements normalized to cell number (Figure 1C-D). Because mitochondrial content changes substantially during pregnancy and lactation, mitolysosome burden alone does not reflect the relative extent of mitochondrial turnover. We therefore normalized mitolysosome burden to mitochondrial content to estimate the relative delivery of mitochondria to lysosomes across the developmental time course (Figure 1E). This analysis revealed a reduction in relative mitochondrial delivery to lysosomes at late pregnancy, followed by increased mitochondrial content and a rebound in mitochondrial turnover as the gland transitioned into lactation.

### Differentiation is accompanied by increased respiratory-chain activity and supercomplex abundance in wild-type HC11 cells

We next asked whether the developmental interval associated with increased mitochondrial turnover was accompanied by a change in mitochondrial function during MEC differentiation. To establish the lactogenic differentiation time course in HC11 cells, we examined undifferentiated cells and cells collected at 8, 24, and 48 h after induction of differentiation. Expression of the milk protein gene *Csn2* (casein beta) increased across the time course, with the highest expression detected at 48 h, confirming progressive acquisition of the lactogenic phenotype (Figure 2A). We then measured oxygen consumption rate (OCR) and extracellular acidification rate (ECAR) using the Seahorse mitochondrial stress test to define changes in cellular energetics during differentiation. Consistent with our previous studies of HC11 differentiation [28] undifferentiated cells exhibited a relatively quiescent energetic phenotype, whereas differentiation was associated with increased mitochondrial respiration and a progressively more energetic state, with the most pronounced shift evident by 48 h (Figure 2B). Thus, lactogenic differentiation is accompanied by substantial remodeling of cellular bioenergetics.

**Figure 2.**
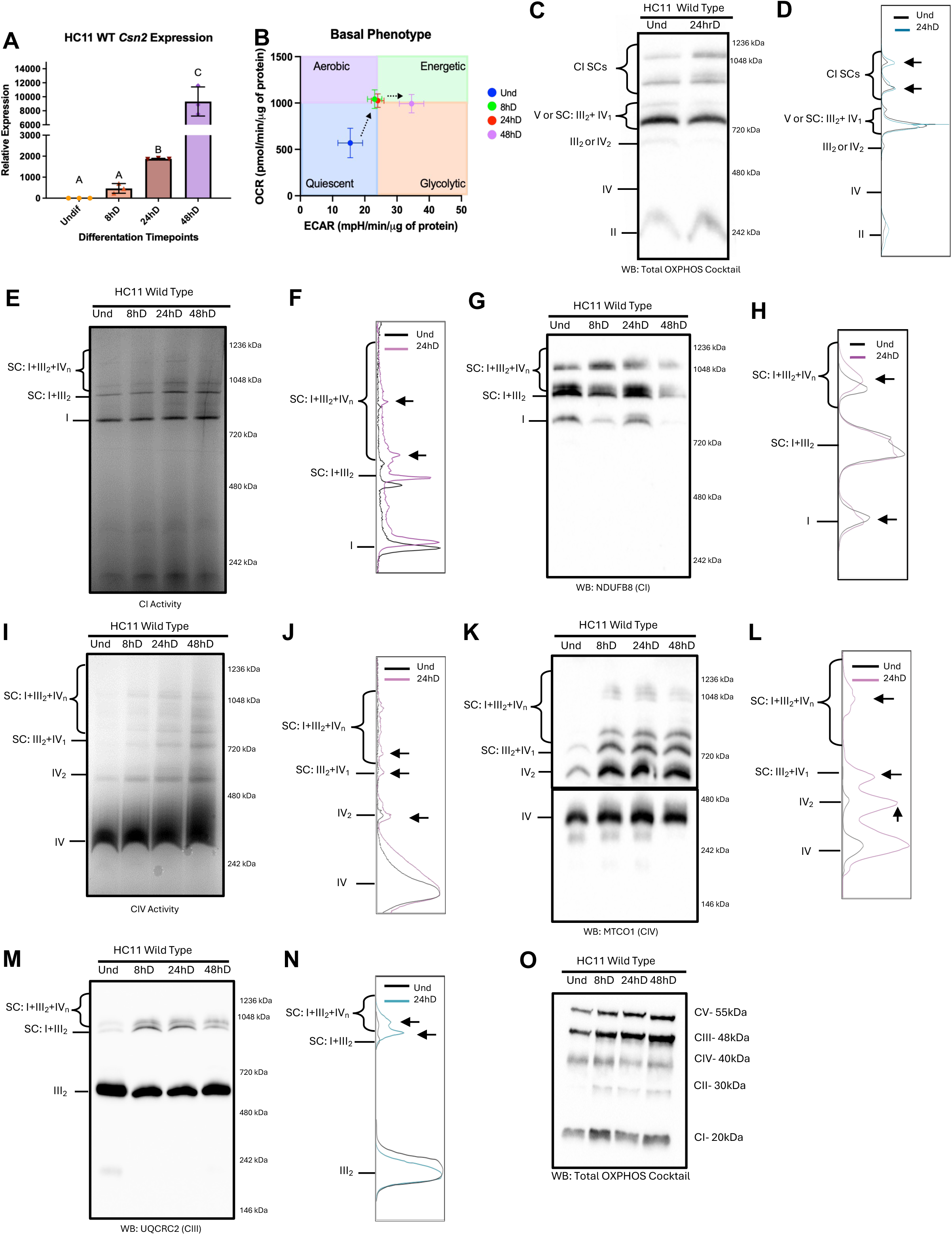
Lactogenic differentiation of HC11 cells corresponds with changes in mitochondrial function and MRC SC formation. (**A**) qPCR of HC11 cells at undifferentiated (Undiff), 8-, 24-, and 48-hours of differentiation (8, 24, and 48hD) for relative expression of milk protein beta casein (*CSN2*). (**B**) Basal energy phenotype comparison of OCRs and extracellular acidification rates (ECARs) in differentiating HC11 cells further demonstrating a dynamic metabolic transition (**C-D**) BN-PAGE analysis of HC11 Wild Type mitochondria at Undifferentiated (Und) and 24hD timepoints (**C**) and densitometry spectra visualizing differences in protein bands for total OXPHOS cocktail (NDUFB8 (CI), SHDB (CII), MTCO1 (CIV), UQCRC2 (CIII), and ATP5A (CV)) **(D)** Arrows indicate particular regions of comparison. (**E-N**) In-Gel activity (**E and I**) and BN-PAGE analysis (**G, K, and M**) of HC11 mitochondria across differentiation timepoints with densitometry analysis highlighting Und and 24hD for Complex 1 (**E-H**), Complex IV (**I-L**), and Complex III (**M-N**). (**O**) Denaturing western of wild type mitochondria probing for total OXPHOS cocktail across differentiation time points. All data are expressed as the mean ± SD, n=3 independent experiments with biological and technical triplicates. For all variables with the same letter, the difference between the means is not statistically significant. Significance was calculated using multiple two-tailed unpaired or Welch’s *t*-tests. *P* < 0.05 was considered statistically significant.

We next determined whether this functional transition was associated with reorganization of the mitochondrial respiratory chain (MRC). Mitochondria were isolated from wild-type HC11 cells across the differentiation time course and analyzed using blue native PAGE (BN-PAGE) and native in-gel activity assays. These approaches preserve higher-order respiratory assemblies, allowing us to assess both the abundance and activity of individual respiratory complexes and supercomplex-containing species.

As an initial assessment of respiratory-chain organization, BN-PAGE followed by immunoblotting with an OXPHOS antibody cocktail was used to compare undifferentiated and 24 h differentiated HC11 cells. Differentiated cells displayed greater signal within higher-molecular-weight regions corresponding to respiratory supercomplexes (Figure 2C). Densitometry profiles further demonstrated enrichment of these higher-order species at 24 h, including the region containing Complex I-associated supercomplexes (Figure 2D).

We next examined individual respiratory complexes across the differentiation time course. Complex I (CI) in-gel activity showed relatively little change in activity associated with monomeric CI; in contrast, NADH oxidation increased within higher-molecular-weight regions corresponding to CI-containing supercomplexes, including SC I+III_2_+IV_n_ and I+III_2_, at 24 and 48 h of differentiation (Figure 2E-F). Consistent with these activity measurements, BN-PAGE followed by immunoblotting for NDUFB8 (NADH:ubiquinone oxidoreductase subunit B8) demonstrated increased incorporation of CI into higher-order assemblies during differentiation, with the greatest abundance detected at 24 h, whereas monomeric CI remained comparatively stable (Figure 2G-H).

A similar pattern was observed for Complex IV (CIV). CIV in-gel activity increased modestly within monomeric and dimeric CIV species, but a more pronounced increase was detected within higher-molecular-weight regions corresponding to CIV-containing assemblies, including III_2_+IV_1_ and I+III_2_+IV_n_, at 24 and 48 h (Figure 2I-J). BN-PAGE followed by MTCO1 immunoblotting similarly demonstrated increased abundance of CIV-containing supercomplexes at 24 h, together with changes in monomeric and dimeric CIV species (Figure 2K-L). Analysis of Complex III (CIII) using UQCRC2 (ubiquinol-cytochrome c reductase core protein 2) immunoblotting further showed increased abundance of CIII-containing higher-order assemblies, including III_2_+IV_1_ and I+III_2_+IV_n_, at 24 h (Figure 2M-N).

To determine whether these changes reflected a general increase in respiratory-chain protein abundance, we performed denaturing immunoblotting for individual MRC complexes across the same differentiation time course (Figure 2O). The marked increase in higher-order respiratory assemblies was not accompanied by a comparable increase in the abundance of most individual respiratory-chain complexes, indicating that differentiation predominantly alters the organization of the respiratory machinery rather than simply increasing its abundance. Together, these data demonstrate that lactogenic differentiation is accompanied by remodeling of respiratory-chain architecture, characterized by increased assembly and activity of CI-, CIII-, and CIV-containing supercomplexes during acquisition of the differentiated state.

### Parkin-dependent mitophagy is required for differentiation-associated respiratory-chain remodeling

To test whether respiratory-chain supercomplex remodeling occurs in conjunction with programmed mitochondrial turnover, we disrupted mitophagy by depleting PRKN (parkin RBR E3 ubiquitin protein ligase) in HC11 cells. Because PRKN is required for mitochondrial turnover and lactogenic differentiation in this model [28], we first confirmed impaired differentiation by the failure of *shPrkn* cells to induce *Csn2* at 24 h of differentiation (Figure 3A). Our previous work has shown that PRKN depletion also prevents acquisition of the energetic phenotype associated with HC11 differentiation [28]. Together, these findings established a model in which programmed mitochondrial turnover is disrupted and allowed us to determine whether PRKN-dependent mitophagy is required for respiratory-chain remodeling.

**Figure 3.**
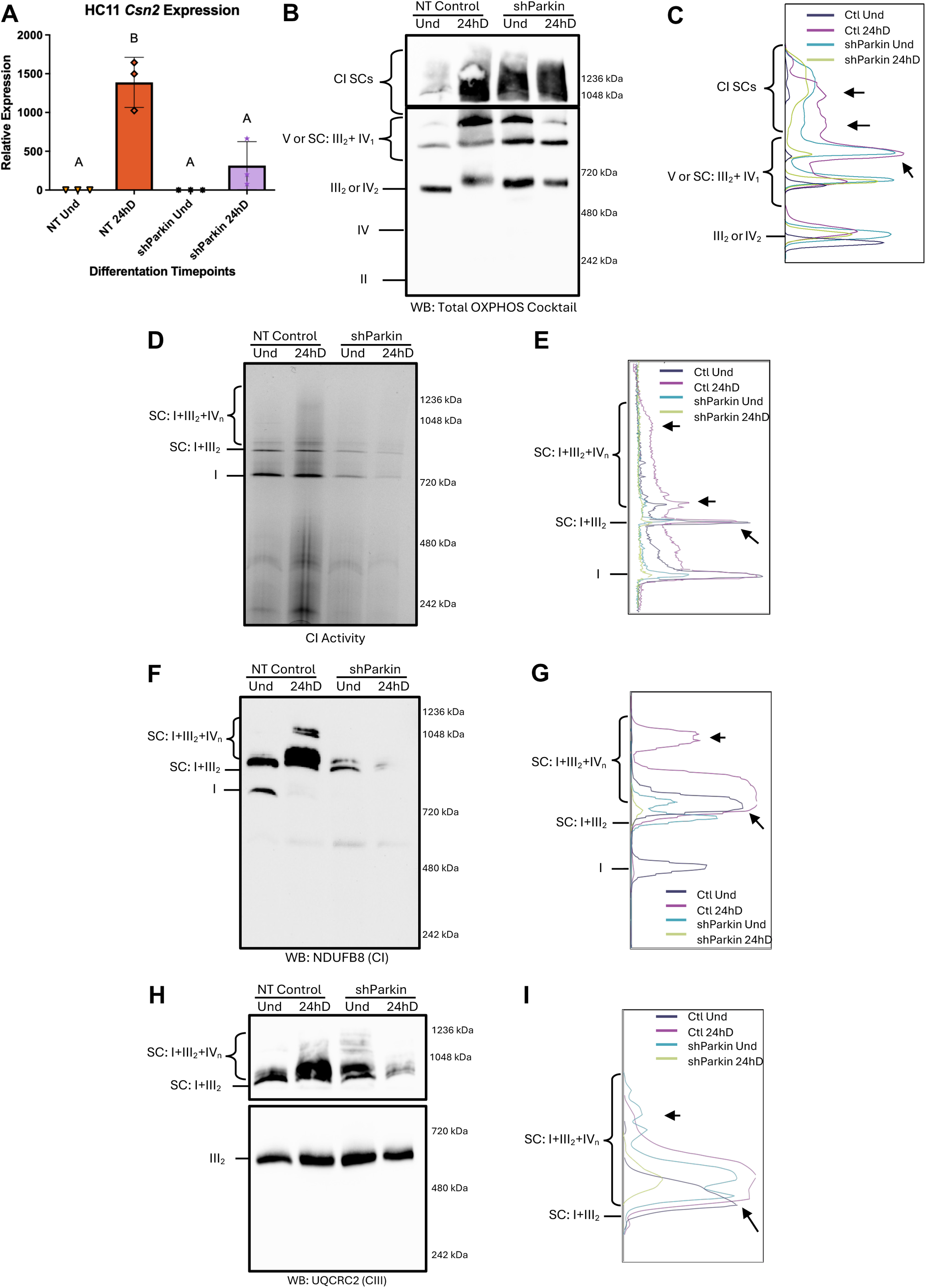
Loss of Parkin-mediated mitophagy and differentiation in HC11 cells corresponds with loss of MRC SC formation. (**A**) qPCR of Non-targeting control (NT, NT control, Ctl) and *Parkin* knockdown (shParkin) HC11 cells at Und and 24hD for relative expression of milk protein beta casein (*CSN2*). (**B-C**) BN-PAGE analysis of NT control and *shParkin* mitochondria at Undifferentiated (Und) and 24hD timepoints (**B**) and densitometry spectra visualizing differences in protein bands for total OXPHOS cocktail (**C**) Arrows indicate particular regions of comparison. (**D-E**) In-Gel activity (**D**) and densitometry spectra (**E**) measuring CI activity in NT control and *shParkin* Und and 24hD mitochondria (**F-I**) BN-PAGE analysis (**F and H**) of NT control and *shParkin* mitochondria at Und and 24hD with densitometry (**G and I**) analysis for Complex 1 (**F-G**), and Complex III (**H-I**). All data are expressed as the mean ± SD, n=3 independent experiments with biological and technical triplicates. For all variables with the same letter, the difference between the means is not statistically significant. Significance was calculated using multiple two-tailed unpaired or Welch’s *t*-tests. *P* < 0.05 was considered statistically significant.

We next isolated mitochondria from non-targeting control and *shPrkn* cells and assessed respiratory-chain organization and activity using native assays. BN-PAGE followed by immunoblotting with an OXPHOS antibody cocktail showed the expected increase in higher-molecular-weight respiratory assemblies at 24 h of differentiation in control cells. In contrast, PRKN-depleted cells failed to acquire this differentiation-associated supercomplex profile (Figure 3B-C). Consistent with this structural defect, Complex I in-gel activity increased within the supercomplex region in control cells at 24 h, whereas this increase was blunted following PRKN depletion. Supercomplex-associated and monomeric CI activity remained reduced in *shPrkn* cells relative to differentiated controls (Figure 3D-E). These findings indicate that PRKN-dependent mitochondrial turnover is required for the normal increase in respiratory-chain activity that accompanies differentiation.

To determine whether this functional defect reflected altered respiratory-chain assembly, we next examined CI- and CIII-containing species by BN-PAGE and immunoblotting. Control cells showed the expected increase in CI-containing supercomplexes at 24 h of differentiation, whereas *shPrkn* cells retained a respiratory-chain profile more similar to the undifferentiated state, with reduced abundance of higher-order CI-containing assemblies (Figure 3F-G). A similar pattern was observed for CIII-containing species, which failed to undergo the differentiation-associated redistribution into higher-order assemblies following PRKN depletion (Figure 3H-I). Together, these data demonstrate that disruption of PRKN-dependent mitochondrial turnover prevents acquisition of the differentiation-associated respiratory-chain architecture, linking programmed mitophagy to remodeling of the mitochondrial respiratory machinery.

### SIM2 alters the timing and magnitude of developmental mitophagy in vivo

Because SIM2 previously emerged as a regulator of mitochondrial turnover in MEC differentiation [18,29,41], we next asked whether SIM2 regulates the developmental timing of mitophagy in vivo. We have previously shown that mammary-specific overexpression of *Sim2s* (MMTV-*Sim2s referred to as Sim2s OE*) increases pup weight during lactation, whereas conditional deletion of *Sim2* using Wap-*Cre* reduces pup weight, consistent with altered mammary epithelial function and milk production [42] To determine whether these phenotypes were associated with changes in mitochondrial turnover, we crossed *Mito-QC* reporter mice with either *Sim2s OE* mice to examine SIM2 gain of function (*Mito-QCxSim2s OE*) or Wap-*Cre*;*Sim2*^fl/fl^ mice (*Mito-QCxSim2^fl/fl^*) to examine conditional loss of function during late pregnancy and lactation (Figure 4A).

**Figure 4.**
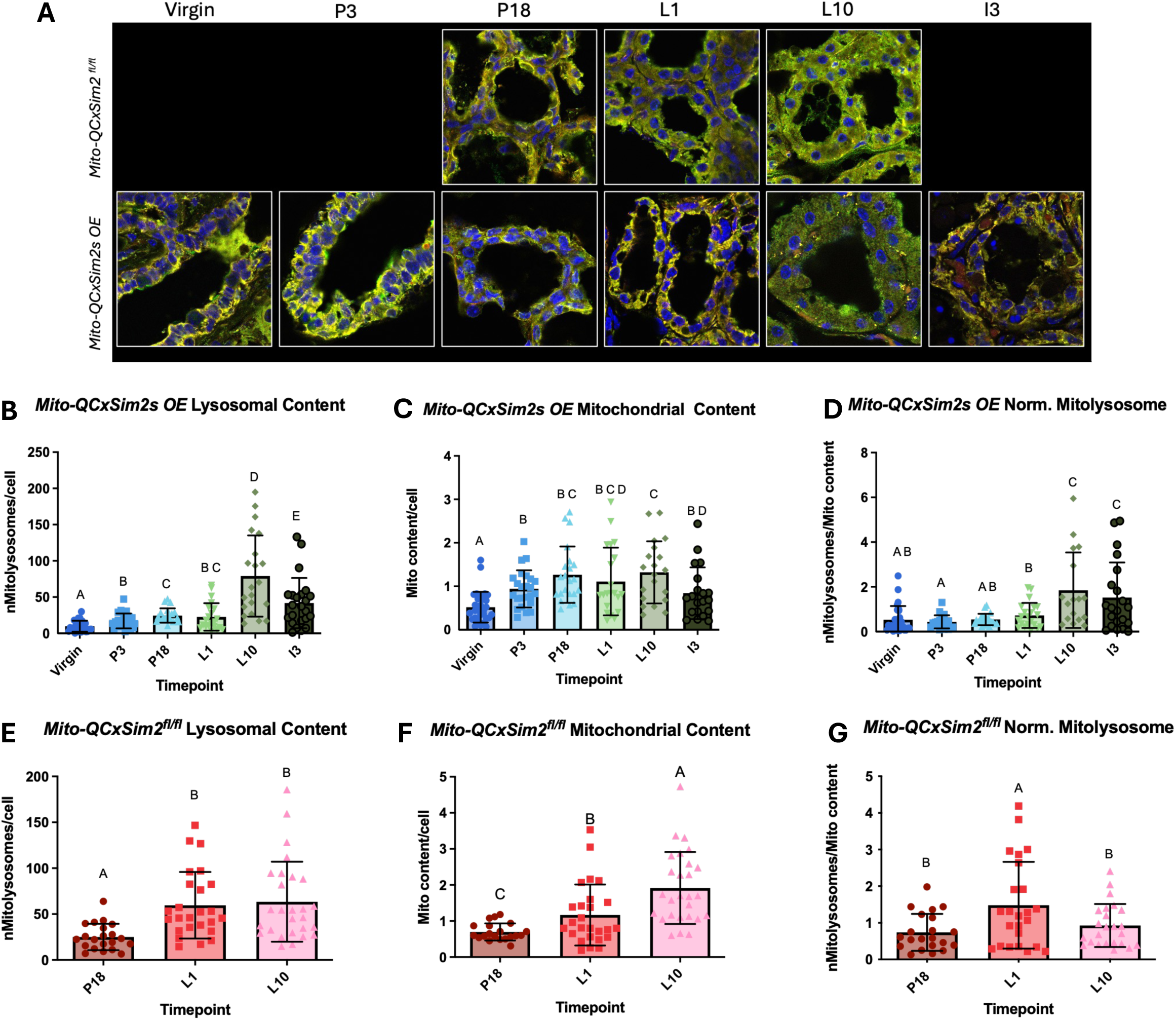
*Sim2* expression alters temporal mitolysosome formation in mouse mammary tissue (**A**) Representative images of *Mito-QCxSim2^fl/fl^ and Mito-QCxSim2s OE* cross mammary glands taken at virgin (in estrus), pregnancy days 3 (P3) and 18 (P18), lactation days 1 (L1) and 10 (L10), and involution day 3 (I3) and stained for immunofluorescence with GFP and mCherry antibodies. *Mito-QCxSim2^fl/fl^* glands were taken only at P18, L1, and L10 due to the temporal control of the conditional knockout. Images were taken using 100x objective. Scale bars 10uM. All timepoints have *n* = 4–6 mice with 3-6 images per mouse. Quantification of mitolysosome number (mCherry^+^GFP^−^ puncta), mitochondrial content (mean GFP intensity), mitolysosomes controlled for mitochondrial content, all normalized to cell count for *Mito-QCxSim2s OE* (**B-D**) and *Mito-QCxSim2^fl/fl^* (**E-G**) time courses. All data are expressed as the mean ± SD. Dots represent individual images. For all variables with the same letter, the difference between the means is not statistically significant. Significance was calculated using multiple two-tailed unpaired or Welch’s *t*-tests. *P* < 0.05 was considered statistically significant.

Quantification of these developmental time courses demonstrated that SIM2 altered the temporal pattern of mitochondrial turnover rather than producing a uniform change in mitochondrial content. In *Mito-QCxSim2s OE* glands, the late-pregnancy reduction in mitochondrial delivery to lysosomes observed in control *Mito-QC* tissue was followed by a more sustained increase across lactation, extending from L1 through L10 (Figure 4B-D). In contrast, conditional loss of *Sim2* altered this pattern, with increased mitolysosome burden at L1 followed by reduced relative mitochondrial delivery to lysosomes at L10 (Figure 4E-G). Thus, both increased and reduced SIM2 dosage disrupted the normal developmental pattern of mitochondrial turnover, identifying SIM2 as a regulator of the timing and persistence of mitochondrial renewal during the pregnancy-to-lactation transition.

Because SIM2 altered developmental mitochondrial turnover at the whole-gland level, we next asked whether these changes differed between mammary epithelial lineages. *Mito-QC* tissues were co-stained for E-cadherin (ECAD) and keratin 14 (K14) to distinguish luminal and basal epithelial populations, respectively (Figure 5A-B). In virgin control glands, K14-positive basal cells exhibited lower mitochondrial delivery to lysosomes than ECAD-positive luminal cells after normalization to mitochondrial content, despite no significant differences in overall mitolysosome burden or mitochondrial content between the two populations (Figure 5C-E). This lineage-specific difference is consistent with previous reports that luminal mammary epithelial cells contain greater abundance of oxidative phosphorylation machinery than basal cells [43,44] suggesting that mitochondrial turnover may differ according to the metabolic requirements of these populations.

**Figure 5.**
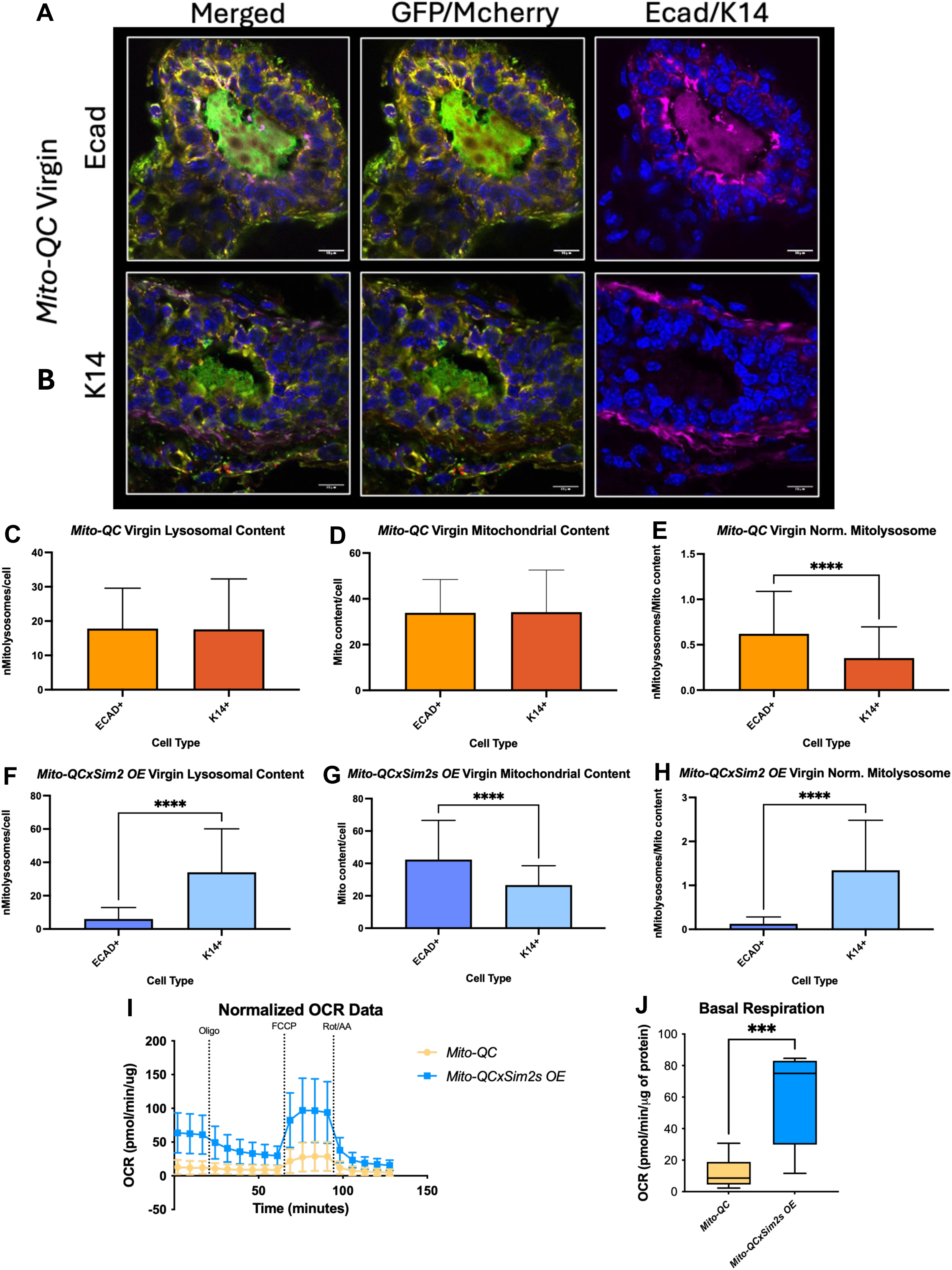
Mammary epithelial cell lineage impacts mitolysosome formation (**A-B**) Representative images of *Mito-QC* and *Mito-QCxSim2s OE* cross mammary glands taken at virgin (in estrus) and stained for immunofluorescence with GFP, mCherry, and ECAD (**A**) and K14 (**B**) antibodies. Images were taken using 100x objective and presented as merged, GFP+/mCherry+ merged, and K14 or ECAD/DAPI merged images. Scale bars 10uM. All timepoints have *n* = 4–6 mice with 3-6 images per mouse. Quantification of mitolysosome number (mCherry^+^GFP^−^ puncta), mitochondrial content (mean GFP intensity), mitolysosomes controlled for mitochondrial content, all normalized to cell count for *Mito-QC* **(C-E)** and *Mito-QCxSim2s OE* (**F-G**). (**I-J**) Respiration measurements from primary organoids collected from *Mito-QC* and *Mito-QCxSim2s OE* virgin glands represented by oxygen consumption over time (**I**) and basal respiration (**J**). N=2 with >7 technical replicates each. All data are expressed as the mean ± SD. ****P<0.0001 ***P<0.001, significance was calculated using a two-tailed Welch’s *t*-test.

SIM2 overexpression altered this lineage-associated pattern. In virgin *Mito-QCxSim2s OE* glands, K14-positive basal cells exhibited greater relative mitochondrial delivery to lysosomes than ECAD-positive luminal cells, reversing the relationship observed in control tissue (Figure 5F-H). Because MMTV-*Sim2s* mice exhibit precocious features of mammary epithelial differentiation [19,30], we next asked whether this altered pattern of mitochondrial turnover was accompanied by a change in mitochondrial function. Primary MECs isolated from virgin *Mito-QCxSim2s OE* glands displayed higher basal OCR than cells isolated from virgin *Mito-QC* glands (Figure 5I-J). Together, these findings indicate that SIM2 alters both lineage-associated mitochondrial turnover and mitochondrial respiratory activity prior to lactation.

### SIM2 associates with respiratory chain supercomplexes and supports their activity and assembly during differentiation

We previously showed in breast cancer cell models that mitochondrial SIM2s promotes respiratory-chain supercomplex assembly and enhances oxidative phosphorylation. [33]. We therefore asked whether SIM2 similarly regulates respiratory-chain organization during mammary epithelial differentiation. SIM2 knockdown in HC11 cells impaired induction of *Csn2* at 24 h of differentiation, confirming the expected loss of SIM2-dependent lactogenic differentiation (Figure 6A). Our previous studies have shown that loss of *Sim2* also prevents HC11 cells from acquiring the energetic phenotype associated with differentiation, supporting a role for SIM2 in mitochondrial adaptation during this transition [29].

**Figure 6.**
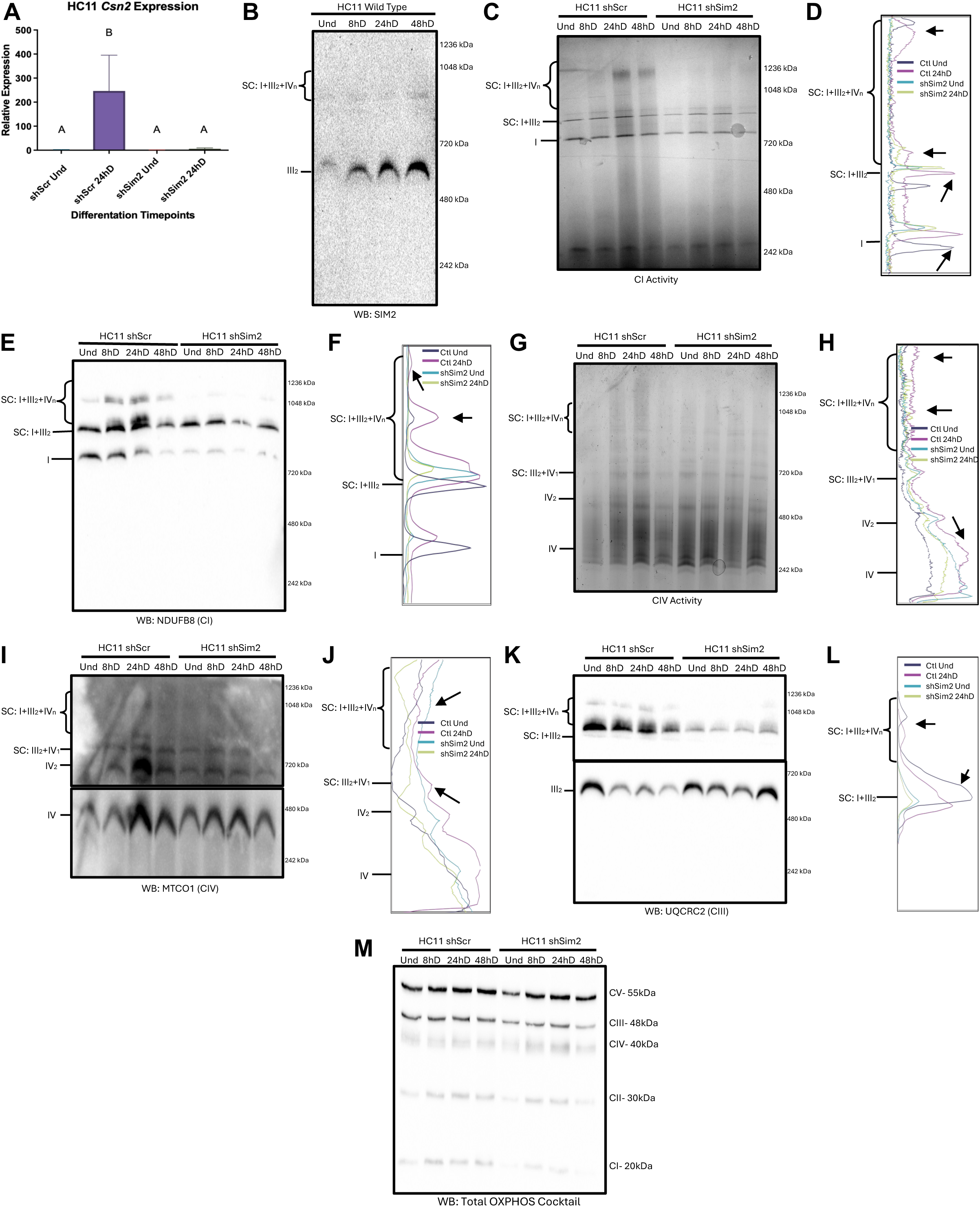
Loss of *Sim2* in HC11 cells corresponds with loss of differentiation-associated MRC SC formation. (**A**) qPCR of scramble control (shScr, Ctl) and *Sim2* knockdown (shSim2) HC11 cells at Und and 24hD for relative expression of milk protein beta casein (CSN2). (**B**) BN-PAGE analysis of wild type HC11 mitochondria across differentiation time points probing for Sim2 (**C-L**) In-Gel activity (**C and G**) and BN-PAGE analysis of *shScr* and *shSim2* mitochondria across differentiation timepoints (**E, I, and K**) and densitometry spectra (**D, F, H, J and L**) visualizing differences in activity/protein bands between *shScr* (Ctl) and *shSim2* Und and 24hD for CI (**C-F**), CIV (**G-J**) and CIII (**K-L**). Arrows on densitometry plots indicate particular regions of comparison. (**M**) Denaturing western of wild type mitochondria probing for total OXPHOS cocktail across differentiation time points. All data are expressed as the mean ± SD, n=3 independent experiments with biological and technical triplicates. For all variables with the same letter, the difference between the means is not statistically significant. Significance was calculated using multiple two-tailed unpaired or Welch’s *t*-tests. *P* < 0.05 was considered statistically significant.

Mitochondria isolated from wild-type HC11 cells across differentiation were analyzed by BN-PAGE followed by immunoblotting for SIM2. SIM2 was readily detected in mitochondrial fractions and increased within the same high-molecular-weight region in which CI-, CIII-, and CIV-containing supercomplexes accumulated, as well as the CIII dimer, with the strongest signal detected at 24 and 48 h of differentiation (Figure 6B). Thus, mitochondrial SIM2 was enriched within the same developmental window and native molecular-weight region in which respiratory-chain remodeling was most pronounced.

To define how SIM2 relates to supercomplex function, we compared mitochondria isolated from control and *shSim2* knockdown cells across differentiation time courses mirroring the wild type timepoints in which function and assembly of supercomplexes occurs at 24 h differentiation. In an in-gel assay, loss of *Sim2* reduced CI activity specifically within the supercomplex region. Whereas control cells increased CI-associated activity at 24 h of differentiation, transitioning into the same supercomplex-active state observed in wild-type cells, *shSim2* cells showed diminished activity in the high-molecular-weight bands despite only modest change in the monomeric region and instead maintained a more immature activity profile (Figure 6C-D). Thus, the loss of *Sim2* phenocopied the failure of respiratory-chain remodeling seen after disrupting mitophagy.

Because *Sim2* loss reduced supercomplex-associated activity, we next examined whether it also altered the structural assembly of these complexes across differentiation. Prior work in MCF7 and SUM159 breast cancer cells showed that SIM2 abundance correlates with the abundance of assembled supercomplexes such that increased SIM2 abundance corresponds with increased assembly and vice versa [33]. We therefore analyzed native assemblies by BN-PAGE across the HC11 differentiation time course. Control cells reproduced the pattern seen in wild-type HC11 mitochondria, with CI-containing supercomplexes increasing during differentiation and reaching maximum abundance at 24 h. In contrast, *shSim2* cells showed a marked loss of high-molecular-weight CI-containing assemblies across differentiation and instead retained a pattern more similar to undifferentiated cells (Figure 6E-F).

CIV activity followed a similar pattern in both in-gel assay and BN-PAGE, with control cells exhibiting differentiation-associated increases in supercomplex-associated cytochrome c oxidation and supercomplex assembly at 24 h that were blunted by *Sim2* knockdown (Figure 6G-J). CIII probing shows again similar patterns. Interestingly, while the control cells demonstrate relative enrichment of the CIII dimer in undifferentiated cells, consistent with redistribution of CIII into higher-order respiratory assemblies during differentiation, the dimer abundance is consistent in the *shSim2* cells even as higher-order assemblies decrease, suggesting the necessity of SIM2 for this reorganization (Figure 6K-L). Indeed, SIM2 probing of control and *Sim2* knockdown cells showed abundance of SIM2 at the molecular weight of the CIII dimer (Figure S3). To determine whether the loss of higher-order assemblies simply reflected reduced abundance of individual respiratory-chain complexes, we performed denaturing immunoblotting for MRC subunits across the differentiation time course (Figure 6M). In control cells, differentiation-associated supercomplex remodeling occurred without broad increases in individual complex abundance, consistent with the pattern observed in wild-type HC11 cells; however, *Sim2* depletion was associated with reduced CI abundance, indicating that *Sim2* loss affects both CI stability or accumulation and its incorporation into higher-order respiratory assemblies. Together, these findings demonstrate that SIM2 is required for the normal acquisition of differentiation-associated respiratory-chain architecture and activity in MECs.

### *Ex vivo* mammary mitochondria recapitulate differentiation-associated remodeling and reveal a requirement for SIM2

We next asked whether the respiratory-chain remodeling observed in HC11 cells could also be detected in primary mammary tissue across development. Because our *Mito-QC* studies identified marked changes in mitochondrial turnover across the transition from late pregnancy to lactation, we isolated mitochondria from mammary glands at late pregnancy (P18), early lactation (L3), and mid-lactation (L10) from control *Mito-QC* and conditional *Mito-QCxSim2^fl/fl^* mice. These time points encompassed the interval of greatest developmental change in mitochondrial turnover and allowed us to determine whether respiratory-chain remodeling occurred in parallel and whether it required SIM2.

Complex I in-gel activity assays supported this model. *Mito-QC* mitochondria showed a lactation-associated increase in CI activity with highest levels appearing in the higher-order regions at L10. In contrast, *Sim2*-deficient mitochondria failed to mount the same increase and instead maintained comparatively low CI-containing supercomplex activity across pregnancy and lactation despite little change in CI monomer activity (Figure 7A-B). BN-PAGE of the same mitochondrial preparations further demonstrated that the ex vivo changes were structural as well as functional. Control tissues exhibited increased abundance of higher-order assemblies during lactation, with the most robust supercomplex signal observed in *Mito-QC* samples at L10 while Sim2-defeicient mitochondria lack supercomplex formation (Figure 7C-D). CIV activity and structure showed a parallel trend: *Mito-QC* samples displayed increased CIV activity and higher-order complex formation at peak lactation, whereas conditional loss of *Sim2* blunted this response (Figure 7E-H). Probing CIII further mirrored this increase with *Mito-QC* CIII-containing supercomplexes increasing during lactation while *Sim2*-deficient mitochondrial failed to establish this reorganization (Figure 7I-J). Thus, primary mammary tissue recapitulated the remodeling observed in HC11 cells and identified SIM2 as a key determinant of lactation-associated supercomplex formation.

**Figure 7.**
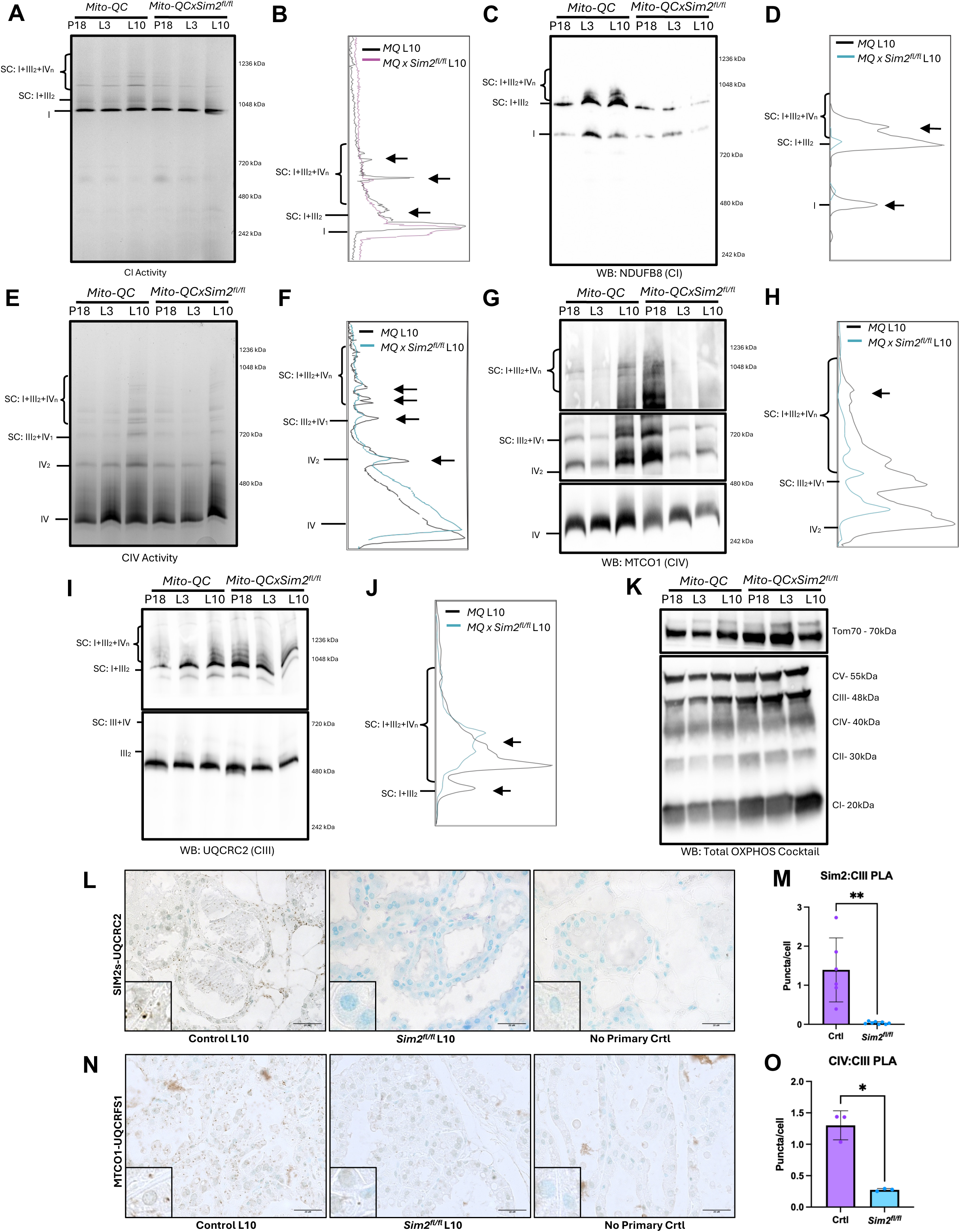
Loss of *Sim2* in primary MEC mitochondria corresponds with loss of differentiation-associated MRC SC formation. (**A-J**) Mitochondria collected from P18, L3 and L10 glands of *Mito-QC* and *MitoQCxSim2^fl/fl^* mice were run with BN-PAGE analysis followed by densitometry for CI (**A-D**), CIV (**E-H**) and CIII (**I-J**). Densitometry analysis of *Mito-QC* (MQ) and *Mito-QCxSim2^fl/fl^* (MQxSim2^fl/fl^) L10 glands (**B, D, F, H and J**) was used to visualize differences in activity/protein bands between genotypes L10 as a representative of peak lactation. Arrows on densitometry plots indicate particular regions of comparison. (**K**) Denaturing western of *Mito-QC* and *Mito-QCxSim2^fl/fl^* mitochondria probing for total OXPHOS cocktail across developmental time points. (**L-O**) Brightfield PLA was performed on L10 glands of *Wap^Cre/+^;mTmG* control mice (Control/Crtl) and *Sim2^fl/fl^* mice (*Sim2^fl/fl^*) probing for proximity of Sim2 and UQCRC2 (**L-M**) and MTCO1 and UQCRFS1 (**N-O**). Images were taken at 63x, scale bar 20uM. N=3 with 3 images per tissue per experiment. Quantification was completed using the cell counter plugin on ImageJ by hand representing average punctate per image normalized for cell count. All data are expressed as the mean ± SD. Significance was calculated using a two-tailed Welch’s *t*-test. *P < 0.05, **P < 0.01

To determine if the *ex vivo* model recapitulated the impacts of *Sim2* loss on relative abundance of individual complexes, we ran denaturing westerns on the mitochondrial samples obtained from the mammary glands (Figure 7K). In *Mito-QC* control mitochondria, the onset of lactation showed no change in the level of individual complexes, mirroring the trends seen in HC11 cells; however, unlike *shSIM2* cells, *Sim2*-deficient mitochondria also showed no change in complex abundance during lactation. This data indicates that changes seen in higher-order complex formation within an *ex vivo* system are not dependent on changes in individual complex abundances.

We have shown previously that SIM2 interacts directly with CIII subunit UQCRC2 via PLA and co-immunoprecipitation in MCF7 cells [33]. Our BN-PAGE results in HC11 cells suggest an interaction with SIM2 and UCQCR2 in MECs *in vitro.* To determine whether SIM2 localized in proximity to respiratory-chain components *in vivo*, we performed brightfield proximity ligation assays (PLA) in lactation day 10 mammary glands of Sim2^fl/fl^ and control mice. Control tissue displayed abundant proximity signal between SIM2 and the UQCRC2, whereas this signal was significantly reduced in *Sim2*-deficient glands (Figure 7L-M). These findings support a close spatial association between SIM2 and Complex III in differentiated mammary epithelium and place SIM2 near respiratory-chain components at the developmental stage when supercomplex remodeling is most pronounced.

We further examined whether proximity between respiratory-chain complexes was altered by loss of SIM2. Brightfield PLA for the CIII subunit UQCRFS1 (ubiquinol-cytochrome c reductase, Rieske iron-sulfur polypeptide 1) and the CIV subunit MTCO1 revealed abundant proximity signal in L10 control mammary epithelium, whereas this signal was significantly reduced in *Sim2*-deficient glands (Figure 7N-O). Together with the native activity and assembly data, these findings support a role for SIM2 in establishing or maintaining higher-order respiratory-chain organization during lactation.

## Discussion

The onset of lactation requires mammary epithelial cells (MECs) to rapidly increase mitochondrial output while simultaneously remodeling the mitochondrial population through programmed turnover. Our findings suggest that this turnover is not simply a response to mitochondrial damage or excess mitochondrial mass. Instead, PRKN-dependent mitophagy is coordinated with SIM2s-dependent reorganization of the mitochondrial respiratory chain (MRC) into active higher-order assemblies during MEC differentiation. In HC11 cells, PRKN-dependent mitochondrial turnover was required for the differentiation-associated increase in the abundance and activity of complex I (CI)-, complex III (CIII)-, and complex IV (CIV)-containing respiratory supercomplexes (SCs). SIM2s regulated mitochondrial delivery to lysosomes in vivo as well as SC assembly and activity in vitro and in mammary tissue. Together, these findings support a model in which programmed mitophagy removes a mitochondrial population that is no longer appropriate for the emerging differentiated state, thereby permitting acquisition of a respiratory-chain architecture better suited to the sustained biosynthetic and secretory demands of lactation [8]. Thus, these studies begin to define what programmed mitophagy builds and identify respiratory-chain architecture as a functional consequence of developmental mitochondrial turnover.

The *Mito-QC* studies extend our previous in vitro observations by mapping developmental changes in mitochondrial delivery to lysosomes in the mammary gland. *Mito-QC* did not reveal a simple progressive increase in mitolysosome burden from pregnancy through lactation. Rather, relative mitochondrial delivery to lysosomes decreased during late pregnancy and was followed by renewed mitochondrial turnover as mitochondrial content increased during lactation. This pattern is more consistent with temporally regulated renewal within an expanding mitochondrial population than with bulk mitochondrial elimination. Overexpression of *Sim2s* broadened and prolonged the lactation-associated changes in mitolysosome burden, whereas conditional loss of *Sim2* altered the early-lactation response and failed to maintain the lactation-associated pattern at L10. Importantly, *Mito-QC* provides a static measure of mitochondrial delivery to an acidic compartment and should not be interpreted as a direct measurement of mitophagic flux. Nevertheless, the contrasting *Sim2* models suggest that SIM2s contributes to the timing and persistence of mitochondrial delivery to lysosomes, consistent with a role in developmental mitochondrial turnover. The lineage-specific analysis further indicates that this process is not uniform across the mammary epithelium. In the virgin gland, ECAD-positive luminal cells showed greater relative mitochondrial delivery to lysosomes than K14-positive basal cells despite similar mitochondrial content and mitolysosome burden. This difference is consistent with previous reports that luminal mammary epithelial cells contain greater abundance of oxidative phosphorylation machinery than basal cells [52,53]. *Sim2s* overexpression reversed this relationship and was accompanied by increased basal respiration in primary MECs, suggesting that SIM2s can alter mitochondrial turnover and respiratory function before the onset of lactation.

Previous work in our laboratory demonstrated that PRKN is recruited to mitochondria during MEC differentiation without a detectable loss of membrane potential and that SIM2s promotes this mitochondrial localization [28,29]. These findings suggested that programmed mitophagy in MECs is initiated by developmental signals rather than by the canonical response to mitochondrial depolarization. In the present study, *shPrkn* cells failed to acquire the differentiation-associated increase in supercomplex abundance and CI activity and instead retained an MRC profile more similar to undifferentiated cells. Similar mitochondrial replacement occurs during myogenic and cardiac differentiation, where clearance of the pre-differentiated mitochondrial population is required for metabolic maturation [12,13]. Loss of *Sim2* resulted in a similar respiratory phenotype, reducing CI-, CIII-, and CIV-containing higher-order assemblies and their associated activity. Together with our previous finding that SIM2s promotes PRKN recruitment, these results position SIM2s as an important regulator of PRKN-dependent mitochondrial turnover and of the respiratory-chain remodeling that accompanies differentiation. SIM2s was present in the same high-molecular-weight region in which respiratory supercomplexes accumulated during differentiation, was closely associated with the CIII subunit UQCRC2 in mammary epithelium, and was required to maintain proximity between CIII and CIV in vivo. We therefore propose that SIM2s coordinates two linked aspects of mitochondrial maturation during MEC differentiation: PRKN-dependent turnover of the pre-existing mitochondrial population and organization of the respiratory machinery within the population that emerges. Although these findings support a functional connection between mitochondrial turnover and MRC remodeling, loss of either *Prkn* or *Sim2* also impairs differentiation, making it difficult to determine whether failed supercomplex remodeling is a direct consequence of disrupted mitophagy or occurs secondary to failure of the broader differentiation program.

The functional significance of respiratory SCs remains debated. SC formation has been proposed to stabilize individual MRC complexes, influence electron transfer, and limit reactive oxygen species production [34,45,46]; however, SC formation does not necessarily imply substrate channeling or increased catalytic efficiency. The failure of these complexes to form could have energetic consequences that play a role in pathologies where the MRC is dysregulated [47–51]. The MRC is also known to respond to environmental stimuli to adapt to energy demands through interactions with other non-respiratory pathways. Studies have shown that mtFAS and fatty acid oxidation proteins work to stabilize MRC super complexes and stabilize OXPHOS levels through physical interaction of proteins such as acyl-CoA dehydrogenase with MRC complexes and donation of electrons into the electron transport chain [52–57]. In our model, the most prominent increase in higher-molecular-weight respiratory assemblies occurred at 24 h of differentiation and preceded maximal *Csn2* expression. This increase was accompanied by greater CI- and CIV-associated activity in the SC region and was recapitulated in mitochondria isolated from lactating mammary tissue. Importantly, the change in SC abundance was not explained by a broad increase in the abundance of individual MRC complexes. Like a Lego build, higher order complexes are made from joining together existing “brick” subunits rather than increases in subunit numbers (Figure S5). Thus, our data support the idea that lactogenic differentiation involves redistribution of existing respiratory complexes into higher-order structures, aligning with the idea that formation of individual subunits and SC formation are intrinsically linked [34,58–60]. We suggest that this reorganization provides a mitochondrial population better suited to the sustained energetic and biosynthetic requirements of lactation, although direct measurements of electron transfer efficiency, substrate use, and reactive oxygen species production will be necessary to define the functional advantage of these assemblies.

These findings may also have relevance to breast cancer, where mitochondrial plasticity associated with normal mammary differentiation can be repurposed during tumor progression. We previously showed that re-expression of *SIM2s* in SUM159 breast cancer cells increased respiratory SC formation and OXPHOS while suppressing proliferation [33]; however, breast cancer cells may uncouple mitochondrial remodeling from differentiation and use the same machinery to adapt to stress. MANF-mediated activation of PRKN-dependent mitophagy supports breast cancer cell survival during glucose starvation and promotes fatty acid oxidation, tumor growth, and metastasis [61]. Conversely, loss of BNIP3-dependent mitophagy in the MMTV-*PyMT* model causes accumulation of dysfunctional mitochondria, increased mitochondrial ROS-HIF1A signaling and glycolysis, and accelerated tumor progression [62]. These apparently opposing outcomes emphasize that the consequence of mitochondrial turnover depends on the cell state in which it occurs. Breast tumors may therefore hijack mitochondrial plasticity normally used during mammary differentiation while uncoupling organelle remodeling from the differentiated state it normally supports [63,64].

Several limitations and mechanistic questions remain. *Mito-QC* cannot distinguish whether changes in mitolysosome burden reflect altered mitochondrial delivery, lysosomal degradation, or mitolysosome residence time, and BN-PAGE and in-gel activity assays do not define the complete composition or intact-cell function of individual MRC assemblies. Differences between the HC11 and conditional mouse models may also reflect incomplete or heterogeneous *Sim2* deletion within the mammary epithelium. Future studies combining pulse-chase labeling with *Mito-QC* and quantitative complexome analysis should determine whether an older mitochondrial population is selectively removed and replaced by mitochondria enriched for specific SC configurations and whether SIM2s also regulates mitochondrial biogenesis or cristae organization. More broadly, programmed mitophagy may represent one component of a coordinated selective-autophagy program that helps establish the differentiated state. Mitophagy supports oxidative transitions in myoblasts, cardiomyocytes, and MECs but a glycolytic transition in retinal ganglion cells [12–14], suggesting that its conserved function may be to replace organelles no longer suited to the emerging cell state rather than enforce a specific metabolic endpoint. This concept may extend beyond mitochondria, as ER-phagy, lipophagy, pexophagy, and ribophagy could work with mitophagy to remodel the organelle network required for lactation. RETREG1/FAM134B-dependent reticulo-mito-phagy provides proof of principle that selective turnover of different organelles can be coordinated [65–67].

In summary, we propose that SIM2s links PRKN-dependent programmed mitophagy to remodeling of the MRC during mammary epithelial differentiation. Together, these processes contribute to the transition from a pregnancy-associated mitochondrial state to one capable of supporting sustained milk synthesis and secretion. Importantly, the mammary gland provides a physiological model in which programmed mitophagy can be examined without chemical depolarization or nutrient deprivation. Our findings support a model in which developmental mitophagy is not simply a mechanism for eliminating damaged mitochondria but a means of enabling acquisition of a mitochondrial state appropriate for differentiated cell function. Future work will be necessary to establish the temporal order of mitochondrial removal, biogenesis, cristae remodeling, and SC assembly and to determine whether similar mechanisms operate in other differentiating tissues. Nevertheless, these results suggest that selective mitochondrial turnover contributes not only to mitochondrial quality control but also to the functional identity of the differentiated cell.

## Materials and Methods

### Mouse lines

All mice were cared for in accordance with the Texas A&M University ethical guidelines and Animal Care and Use Committee. Mice were provided access to food and water ad libitum and were housed under standard 12 h photoperiods. Only female mice were used for this study as it concerned mammary gland development. No method of randomization was used. Three to five female mice were analyzed for each timepoint. Timepoints were chosen to represent of the morphological and functional changes that occur during mammary gland development. Mice collected at virgin timepoints to were staged for collection in estrus. After sacrifice, mammary glands were harvested for immunostaining and mitochondrial isolation. Transgenic mice were bred and genotyped by the Texas A&M Institute for Genomic Medicine. MMTV-*Sim2s* mice have been described previously and were bred in an FVB background [41]. *Sim2^fl/fl^* mice have been described previously and are bred on a mixed 129/SvEv and C57BL/6 background and backcrossed to a C57BL/6 background to produce *Sim2^fl/fl^;Wap^Cre/+^;mTmG* mice (*Sim2^fl/fl^*) and littermate *Wap^Cre/+^;mTmG* (control) mice [29]. Rosa26-mCherry-GFP-FIS1^101-152 (*Mito-QC)* reporter mice were developed by the Ganley lab and generously gifted to the Porter lab [37]. MMTV-*Sim2s* and *Sim2^fl/fl^* mice were then crossed with the *Mito-QC* mice to produce an F1 generation of mcherry/GFP/FIS1xMMTV-*Sim2s* homozygotes (*Mito-QCxSim2s OE*) and mCherry/GFP/FIS1xTom/Cre/Sim2^fl/fl^ homozygotes *(Mito-QCxSim2^fl/fl^*).

### Cell Culture

HC11 cells (ATCC, CRL-3062) and were maintained and differentiated as previously described [28,29,41,68]. Briefly, HC11 cells were grown in RPMI (Gibco, 22400121) supplemented with 10% fetal bovine serum (Sigma), 50 μg/mL gentamycin (Gibco, 15750078), 5 μg/mL bovine insulin (MilliporeSigma, I5500-500MG), and 10 ng/mL recombinant mouse EGF (Gibco, PMG8041). For differentiation timepoints, HC11 cells were grown to confluence, after which growth medium was replaced with priming medium (RPMI supplemented with 10% donor horse serum, 50 μg/mL gentamycin, 5 μg/mL bovine insulin, and 1 μg/mL hydrocortisone [MilliporeSigma, H0888-1G]). After 24 h, priming medium was replaced with differentiation media, which consisted of priming media plus 1 μg/mL ovine prolactin (R&D Systems 682-PL-050). Differentiation media remained on the cells for the duration of the designated timepoints. Differentiation was considered initiated from addition of differentiation medium. Stable *shSim2* and *shParkn* cell lines have been previously validated and described [20,24,28,29,41,69].

### Mitochondrial Isolation

Mitochondria are harvested from HC11 cells as previously described [33,70] after scraping by pelleting cells via centrifugation at 1000rpm at 4°C for 2 minutes. Pelleted cells are then resuspended in 5.5mLs of cold RSB hypo buffer (10 mM NaCl, 1.5 mM MgCl2, 10 mM Tris-HCl, pH 7.5) and kept on ice for 10 minutes to allow cell lysis. Samples are then lysed with the aid of a glass-on-glass Dounce tissue homogenizers using 20 strokes with both the loose and tight pestles. Lysis was halted by the addition of MSH buffer (210 mM mannitol; 70 mM sucrose; 5 mM Tris-HCl; and 1 mM EDTA, pH 7.5). Both RSB and MSH buffers included 1 mM Na3VO4 and 1 mM cOomplete ULTRA Tablets, Mini, EDTA-free EASYPack (Roche, 5892953001) to inhibit phosphatase and protease activity as well as 100 nM MG-132 (Cayman Chemical Company, 13697). The cell suspension was then spun at 600 x g for 5 minutes at 4°C, after which a pellet of nuclear fraction and cellular debris is evident. The supernatant was decanted off and the spin repeated, after which the supernatant was centrifuged at 7000 x g for 10 minutes at 4°C in a slant rotor. After this spin, a pellet of mitochondria was visible, while the cytosolic proteins remain in the supernatant. The supernatant was decanted, and the pellet resuspended in 3 mLs of cold MSH buffer and the centrifugation repeated. Finally, the cell suspension was spun at 10,000 x g and the supernatant discarded, leaving a crude mitochondrial pellet that was resuspended in MSH buffer, assayed using a DC protein assay kit (Bio-Rad, 5000112) and aliquoted to be used to native applications.

Primary mitochondria were isolated from frozen mouse glands flash frozen in liquid nitrogen and stored at −80°C according to protocol modified from Osto *et al.* (2020) [71]. Shortly, previously frozen glands were weighed, thawed in cold PBS and minced before homogenization in 250 uL per 10 mg of tissue in cold MAS buffer (70mM sucrose, 220mM mannitol, 5mM KH_2_PO_4_, 5mM MgCl_2_, 1mM EGTA, and 2mM HEPES) with inhibitors (identical to those listed above for MSH buffer) using a glass-on-glass homogenizer using 20 strokes with both the loose and tight pestles. Homogenates were then centrifuged twice for 5 minutes at 1000 x g in a precooled 4°C centrifuge, discarding the pellet, collecting the supernatant, and carefully aspirating off fat layers after each spin. Supernatants were then centrifuged for 10 mins at 10,000 x g at 4°C in a slant rotor. After this spin, the supernatants were decanted, and the pellets resuspended in 3mLs of cold MAS buffer and the centrifugation repeated twice. Finally, the pellets were resuspended in 1 mL of MAS and spun a final time at 10,000 x g and the supernatant discarded, leaving crude mitochondrial pellets that were resuspended in MAS buffer and handled identically to mitochondria from cultured cells.

All isolations where repeated for a minimum of 3 independent collections as biological replicates.

### Primary organoid isolation

Primary Mammary Epithelial cells were harvested from virgin *Mito-QC* and *Mito-QCxMMTV* cross glands according to protocol adapted from Welm *et al*. (2008) [72]. Shortly, primary MECs were isolated from the inguinal and thoracic mammary glands, pooled from 2-3 mice per genotype, and placed in wash buffer (1xDMEM/F12 [Gibco, 11320082], 5% FBS [Sigma F2442], and 50 µg/mL gentamicin [Gibco, 15750078]) with 2mg/mL Collagenase A (Roche, 11088793001) after dissection with light homogenization with scalpels (VWR,82029-860). Homogenates were then incubated with moderate shaking for 1.5 hours at 37°C. Organoids were pelleted at 600 x g for 10 minutes in a cooled centrifuge, after which supernatants were aspirated and the pellet was treated with DNase I (100 µg/mL DNase I [Roche, 10104159001] in DMEM/F12) and were enriched by 3 rounds of washing and pulse spinning at 450 x g to obtain a pellet of isolated mammary organoids.

Isolated organoids were then plated and run through a Seahorse Organoid Assay 3D Mito Stress Test (Agilent, 103016-100) according to the manufacture protocol with 2-step seeding procedure. Shortly, during the DNase and wash steps of organoid prep, a XF Flex organoid microplate (Agilent, 103865-100) was loaded with 6 uL/well of Cultrex (R&D Systems, 3432-005-01) using cold pipett tips and then incubated at 37°C for 20 minutes to allow the Cultrex to solidify. The organoids were resuspended in 100% Cultrex after their final wash and pulse spin and 4 uL of the suspension loaded into each well of the plate except A1 and D6 which serve as background wells and received 4 uL of plain Cultrex. The plate was incubated again for 20 min at 37°C to allow polymerization after which 250 uL of prewarmed growth media (1x DMEM/F12, 10uL/mL Penicillin-streptomycin [Gibco, 15140122], and 10 uL/mL Insulin-Transferrin-Selenium-Ethanolamine [Gibco, 51500056]) was added to each well. The plate was maintained overnight prior to running the Mito Stress Test. The Mito Stress Test was run as described in this paper for cultured cells, with final concentrations of 20 uM Oligomycin, 15 uM FCCP, and 10 uM Rot/AA after which the assay was normalized to protein content per well.

### Histology

Tissue samples were fixed overnight at 4°C in 4% paraformaldehyde and stored in 70% ethanol at 4°C until processing. Samples were processed, sectioned, and H&E stained by the Texas A&M University College of Veterinary Medicine & Biomedical Science Histology Laboratory. Immunostaining was performed as previously described [28,41]. Briefly, tissue sections were de-paraffinized and rehydrated in graded ethanol washes. Antigen retrieval was performed by incubation in 10 mM sodium citrate under high pressure for 5 min, and peroxidases were blocked in 3% hydrogen peroxide for 6 min. Blocking was performed in 10% horse serum or Mouse on Mouse (M.O.M) blocking diluent (Vector Laboratories, BMK-2202) for 1 h at room temperature, and samples were incubated with primary antibodies overnight at 4°C. Biotinylated secondary antibodies were applied for 1 h at room temperature, and samples were subsequently incubated with avidin-biotin peroxidase using the ABC method (Vector Laboratories, PK-6200). Finally, peroxidase was visualized with the chromogen DAB (3,3’-diaminobenzidine; Vector Laboratories, SK-4100), after which tissues were counterstained with Methyl Green, dehydrated, and coverslips were applied with Permount Mounting Medium (VWR, 100496-552). Images were collected on a Zeiss Axio Imager.Z1 with a 63× plan-apochromat objective.

### Proximity Ligation Assay

Interactions between proteins of interest were analyzed by proximity ligation assay (PLA) using MilliporeSigma Duolink Brightfield PLA technology following the manufacturer’s protocol. Mouse mammary sections from selected genotypes and timepoints were collected and processed as described above. Slides underwent de-paraffinization, rehydration, antigen retrieval, and peroxidase blocking as described [29]. Slides were blocked with M.O.M blocking reagent for 60 min at 37 °C and then incubated in primary antibody for overnight at 4°C. Primary antibodies were made up in M.O.M kit antibody diluent. Samples where no primary antibody was added (incubated in M.O.M diluent) were included as negative controls. After primary antibody incubation, Duolink PLA probes, anti-mouse PLUS (DUO92004) and anti-rabbit MINUS (DUO92002) were added to the slides and incubated at 37°C for 1h. Signal was generated using Duolink Brightfield Detection Reagents (DUO92012) following the manufacturer’s protocol of ligation for 30 min at 37°C and amplification for 120 min at 37°C, followed by detection for 1h at room temperature. After detection, interactions were visualized using DAB and counterstained as described. Finally, slides were dehydrated and coverslips applied with Permount. Images were collected on a Zeiss Axio Imager.Z1 with a 63× plan-apochromat objective.

### Immunofluorescence

Mouse mammary glands from *Mito-QC* and *Mito-QCxSim2s OE* mice were harvested across differentiation timepoints consisting of 10-week virgin (Virgin), pregnancy day 3 (P3), pregnancy day 18 (P18), day 1 of lactation (L1), day 10 of lactation (L10), and after 3 days of forced involution (I3). *Mito-QCxSim2^fl/fl^* mice were collected at P18, L1, and L10 due to the lox system being under control of the whey acidic protein (WAP) promoter which is active only during late pregnancy and throughout lactation. Tissues were fixed at 4°C in PFA for 2 (L10 and I3), 3 (P18, L1), or 6 hours (Virgin and P3) and then washed 3x in cold PBS for 10 minutes. Tissues were then left in a 15% sucrose solution overnight. Mammary glands were embedded in gelatine (MilliporeSigma, G2500-500G) to be used for cryosectioning. Tissues were incubated in warmed 7.5% porcine gelatine for 1 hour prior to embedding. Gelatine infused tissues were placed in molds, flash frozen in 2-methyl butane (MilliporeSigma, M32631-500ML) cooled to - 50° to −60°C with liquid nitrogen and then stored at −80°C.

Tissues were sectioned according to the methods put forth by Yang et. al [73] on a CryoStar NX70 cryostat into 8-micron sections using a tape transfer system and Norland Optical Adhesive 63 (Norland Products, 6301). Shortly, frozen sample blocks were removed from the cassette and mounted onto the specimen chuck using Tissue-Tek O.C.T. (VWR, 25608-930). The block was trimmed by slicing at a thickness of 30 μm until the majority of the tissue was at the face of the block. A segment of tape (Possible Missions, 5030381) was then attached to the trimmed block by placing the adhesive side to the tissue block and firmly rolling it onto the block using a prechilled roller. An 8 μm section was then cut at a slow and placed on a prepared slide with the tissue side facing down. Slides were prepared by adding a small line of optical adhesive to the slide and gently spread to a thin layer using a cell scraper and prechilled. The tissue was lightly pressed into the optical adhesive on the slide to ensure transfer. The slide was then inserted into a UV transilluminator (Accuris, E3100) and then cured with UV light for 30 seconds after which the tape was carefully removed with cold forceps leaving the tissue intact on the slide. The slides were then air-dried overnight before being stored in at 4°C.

Tissue slides were stained for immunofluorescence using antibodies for GFP, mCherry, and other targets according to the concentrations listed in table 1. Briefly, gelatin was removed from slides via immersion in PBS at 37°C for at least 15 min followed by blocking for an hour in PBS working buffer containing 10% BSA (Bovine Serum Albumin; MilliporeSigma, A9647-100G) and Triton-X, 3 10-minute washes in PBS and then incubation in primary antibody at 4°C overnight. After PBS washes the following day, slides are incubated in fluorescent secondaries at 1:1000 dilution for 1 hour at room temp. Finally, each slide was incubated in DAPI (4’,6-diamidino-2-phenylindole, dilactate; Invitrogen, D21490), for nuclear staining for 12 minutes. Slides were then mounted with Fluoro-Gel Mounting Medium with TES Buffer (Electron Microscopy Sciences 17985-30) cover-slipped and sealed at the edges with clear nail polish.

### Blue Native Polyacrylamide Gels (BN-PAGE), and In-Gel Activity Assays

Samples for Blue Native Gels were prepared by resuspending 50 μg of mitochondrial pellets isolated as previously described [33,74] in 1xNativePAGE Sample Buffer (Invitrogen, BN2003), digitonin (4 g/g or 8 g/g) (Invitrogen, BN2006), and water to a final volume of 20 μL and mixed by pipetting. Samples were incubated on ice for 15 min to solubilize the mitochondrial proteins, followed by centrifugation at 4°C for 30 min at 18,000 x g, and the supernatants transferred to new tubes. Lastly, 0.5% G-250 Coomassie dye (Invitrogen, BN2004) was then added to the samples.

BN-PAGE gels were run in an XCell SureLock Mini-Cell (Invitrogen, <u>EI0001</u>) with a NativePAGE Novex 3-12% Bis–Tris gel (Invitrogen, <u>BN1001BOX</u>) as previously described [33]. Shortly, 1xAnode buffer (Invitrogen, BN2007), Dark blue cathode buffer and Light Blue Cathode buffer were mixed according to the NativePAGE Novex Bis–Tris Gel System protocol. After assembly, the wells of the gel were filled with dark blue cathode buffer, followed by samples and 10uL of NativeMark unstained protein standard (Invitrogen, <u>LC0725</u>). The inner chamber of the assembly was filled with Dark Blue Cathode buffer, and the outer chamber was filled with anode buffer until approximately 1/3 full. The gel was run at 4°C for 30 min at 150 V after which the dark blue cathode buffer was replaced with the light blue cathode buffer. The gel was then run for an additional 90 min at 250 V.

After electrophoresis was completed, the proteins were transferred to a membrane using a XCell II Blot Module (Invitrogen, <u>EI9051</u>) onto Sequi-Blot PVDF membranes (BioRad 1620184). Both gel and membrane were incubated in 1x NuPage Transfer Buffer (Invitrogen, <u>NP00061</u>) for 5-15 minutes prior to assembly in the blot module. The blot was transferred at 25V for 1 hour and 15 mins. After the transfer the blot was incubated in 20 mL of 8% acetic acid for 15 min to fix the proteins, after which it was rinsed with DI water and allowed to dry before stripping the background dye via rewetting with 100% methanol and washed again with DI water. Finally, the membrane was blocked for 1 hour at room temp in 5% milk in TBST (Great Value) and then probed and imaged as described for western blots [24,33] with antibodies listed in Table S1.

Samples for in-gel activity assays were prepared by resuspending 30 ug (for complex I) or 100 ug (for complex IV) of isolated mitochondria in 1x NativePAGE Sample Buffer, digitonin (0.4% for complex I in-gel activity, and 1% for complex IV in-gel activity), and water to a final volume of 20 uL and mixed by pipetting. Samples were then prepped identically to BN-PAGE samples described above, without the addition of G-250 dye. The gels were run similarly to the BN-PAGE assembly on a 3-12% Bis-Tris gel, with cathode buffers consisting of Light Blue Cathode In-Gel buffer and Clear Cathode In-Gel buffer mixed as previously described with the addition of DDM (n-dodecyl β-D-maltoside; Invitrogen, BN2005) and sodium deoxycholate (DOC) [33]. After assembly, the wells of the gel were filled with light blue cathode buffer, samples and NativeMark unstained protein standard. The inner chamber was filled with light blue cathode buffer, and the outer chamber was filled with anode buffer until approximately 1/3 full. The gel was initially run at 4°C for 30 min at 100 V after which the light blue cathode buffer was replaced with cathode buffer and the gel run for an additional 4 h at 400 V. After electrophoresis was completed, the gel was treated with staining solutions for either Complex I (25 mg Nitrotetrazolium blue [MilliporeSigma, N6876-100MG], 1mg NADH, 2 mM Tris-HCl) or Complex IV (5mg diaminobenzidine, 50 mM sodium phosphate, 5 mM cytochrome C [MilliporeSigma, C2506-100MG]) overnight. After incubation, the gels were imaged using a ChemiDoc MP Imaging System (Bio–Rad).

All native applications have at least n=2 replicates. Even loading of native gels was determined via parallel western blot immunoblotted for mitochondrial loading controls. Loading blots are available in the supplemental materials.

### Cellular respiration and glycolysis

Cellular respiration was analyzed on a Seahorse XF Flex Analyzer using a V28 plate (Agilent,102342-100). Briefly, cells were seeded at a density of 20,000 per well, growth media was replaced with priming media once cells reached confluence, and the media was again replaced by differentiation media containing prolactin after 24 h, as described. The Seahorse XF Cell Mito Stress Test Kit (Agilent Technologies, 103015–100) was used to measure the oxygen consumption rate (OCR) and the extracellular acidification rate (ECAR), according to the manufacturer’s protocol with two exceptions; oligomycin was used at a final concentration of 2.5 µM, and FCCP was used at a final concentration of 1 µM. Cell number was normalized to the protein content of each well, measured by DC protein assay (Bio-Rad, 5000112). Mean OCR and ECAR values from a minimum of four replicates per group were compared. Quadrant ranges were assigned based on consecutive time points compared by student’s t-test, and quadrants were assigned where the least significant difference between consecutive time points occurred, which indicates a static or change point during the metabolic transition.

### RNA isolation and quantitative real-time PCR (qPCR)

RNA was extracted from cells using High Pure RNA Isolation Kits (Roche, 11828665001) per the manufacturer’s protocols. Reverse transcription was performed with 1 µg total RNA using the iScript cDNA Synthesis Kit (Bio-Rad, 1708891BUN). Subsequent qPCR was performed with 4 µl of cDNA, 6 µM forward and reverse primer mix and SYBR Green Master Mix (Applied Biosystems, A46109) on a CFX384 qPCR (Bio-Rad). Primers for *Csn2 (casein beta*) and *Actb (actin beta*) were synthesized by Integrative DNA Technologies, and the following primer sequences: *Csn2* forward 5’-TGTGCTCCAGGCTAAAGTTCACT-3’, *Csn2* reverse 5’-GGTTTGAGCCTGAGCATATGG-3’, *Actb* forward 5’-GCAACGAGCGGTTCC-3’, and *Actb* reverse 5’-CCCAAGAAGGAAGGCTGGA-3’. The 2−Δ ΔCt method was used to analyze qPCR data, and normalization was performed relative to *Actb*. The standard deviation of the target gene and reference gene Ct values were used to calculate the sum of squares of the standard deviation of each group. This value was used to find the positive and negative errors. The ΔΔCt, positive error, and negative error values were then log-transformed and presented as the fold ± 1 standard deviation. Statistical analyses were performed on the ΔCt values. All experiments had n=3 independent experiments with technical and biological replicates.

### Immunoblotting

Protein was isolated from cells as previously described [24] and lysed in high salt lysis buffer (50 mM HEPES, 500 mM NaCl, 1.5 mM EDTA, 10% glycerol, and 1% Triton X-100 at pH 7.5) containing 1 mM Na3VO4 and 1 mM cOomplete ULTRA Tablets, Mini, EDTA-free EASYPack (Roche). Cell protein concentrations were assessed by DC protein assay (Bio-Rad, 5000112). Equivalent amounts of protein were combined with 6x Laemmli buffer (250 mM Tris-HCl, 8% SDS, 40% glycerol, and 0.4 M dithiothreitol, pH 6.8) and heated at 95°C for 5 min prior to loading on 10% or 12% SDS-PAGE gels. Western blotting was performed as previously described [24,33], and the antibodies used are listed in Table 1. Bands were visualized using ProSignal Pico ECL spray (Prometheus, 20-300S) and digitized on a ChemiDoc MP Imaging System (Bio-Rad) with settings to highlight oversaturated pixels if present.

Native applications were visualized using Fiji (Version 2.0 NIH). In-gel activity assays were preprocessed by removing the background at a rolling ball radius of 500 pixels with a sliding paraboloid and enhancing contrast by .5%. Densitometry spectra of supercomplex regions were overlaid using Adobe Illustrator 2026.

### IF Image Analysis

Slides stained for immunofluorescence were imaged on an Olympus Fluoview FV3000 Confocal Laser Scanning Microscope using the Texas A&M Integrated Microscopy and Imaging Laboratory (IMIL) facilities. Images were taken using a U Apo N 100XTIRF objective. Image analysis was performed on FIJI (NIH). *Mito-QC* and cross images were quantified using the *Mito-QC* ImageJ plugin developed by the Gainly lab [37,39]. In short, merged images were used to create a region of interest (ROI) in which the plug-in would calculate number of mitolysosomes, mitochondrial content, and other values based on user-specified thresholds comparing red (mCherry) and green (GFP) channels. To accurately set thresholds to obtain true quantification of mitochondrial delivery to the lysosome, a “subtraction layer” was obtained via subtracting the green channel from the red channel according to the procedure used by the Boya lab [75] consisting of preprocessing with a background subtraction with a rolling ball radius of 25 pixels and a Gaussian blur filter with a Σ radius of 1. Comparison of the subtraction layer to the corresponding “mitophagy mask” created by the plug-in allowed for tailor thresholding of the plug-in to ensure true quantification. Values obtained from plug-in analysis were then normalized to cell count for each image. In the case of K14+ and Ecad+ analysis, ROIs were determined via the K14/Ecad layer of each image in which each positive cell constitutes an ROI that was then transferred to the merged mCherry/GFP image for analysis for only K14+/Ecad+ cells respectively.

### Statistical analysis

All experiments at minimum were performed with two independent experiments, with biological and technical triplicates where possible. Graphs demonstrate error as ± standard deviation. Box and whisker plots represent the 25th to 75th percentile, and the whiskers represent the minimum and maximum values. The central line indicates the mean of each group. Statistical analyses were performed in GraphPad Prism. ROUT test with Q=1% was used to test for outliers. F tests were conducted to test for significant differences in the variance (significant at *P* < 0.05). When variances were not significantly different, unpaired two-tailed Student’s *t* tests were performed. When variances were different, unpaired two-tailed Welch’s *t* tests were performed*. P* < 0.05 was considered statistically significant.

## Acknowledgements

We would like to thank the Texas A&M University College of Veterinary Medicine & Biomedical Sciences (CVMBS) Histology Laboratory for preparing tissue samples for immunostaining. The authors acknowledge the assistance of the Integrated Microscopy and Imaging Laboratory at the Texas A&M University Naresh K. Vashisht College of Medicine. RRID:SCR_021637. The authors are also grateful to the Texas A&M Institute for Genomic Medicine for production of transgenic mouse lines, maintenance of mouse colonies, and timed matings for the lactation studies and to Dr. Ian G. Ganley for the gift of the Mito-QC mice used in this study. BioRender.com was used to make the models presented in this paper.

## AI Disclosure

No generative AI was used in this research or manuscript preparation process.

## Data Availability Statement

There is no data set associated with this study.

## Abbreviations

BN-PAGE: blue native polyacrylamide gel electrophoresis
CI: complex I
CIII: complex III
CIV: complex IV
CSN2: casein beta
ECAD: E-Cadherin
ECAR: extracellular acidification rate
I: involution day 3
K14: keratin 14
L1: lactation day 1
L10: lactation day 10
MEC: mammary epithelial cell
MRC: mitochondrial respiratory chain
NDUFB8: NADH:ubiquinone oxidoreductase subunit B8
OCR: oxygen consumption rate
P3: pregnancy day 3 P18 pregnancy day 18
PRKN: parkin RBR E3 ubiquitin protein ligase
SC: super complex
SIM2s: single-minded 2s
UQCRC2: ubiquinol-cytochrome c reductase core protein 2
UQCRFS1: ubiquinol-cytochrome c reductase, Rieske iron-sulfur polypeptide 1

