## Supplemental Figures for "SIM2s coordinates programmed mitophagy with respiratory chain supercomplex remodeling during mammary epithelial differentiation"

Table S1. List of Antibodies and concentrations

| Ab | Host | Supplier and cat. number | Application | Concentration |
| --- | --- | --- | --- | --- |
| mCherry | Rabbit | Thermo Scientific, PA5-34974 | IF | 1:100 |
| GFP | Chicken | Aves Labs, GFP-1020 | IF | 1:400 |
| E-Cadherin | Goat | R&D Systems, AF748 | IF | 1:150 |
| K14 | Guinea Pig | OriGene Technologies, BP5009 | IF | 1:250 |
| Tom70 | Rabbit | Proteintech, 14528-1-AP | WB | 1:1000 |
| VDAC | Mouse | Abcam, ab14734 | WB | 1:1000 |
| MTCO1 | Mouse | Abcam, ab14705 | WB, PLA | 1:1000, 1:100 |
| UQCRC2 | Mouse | Abcam, ab14745 | WB, PLA | 1:1000, 1:100 |
| NDUFB8 | Mouse | Abcam, ab110242 | WB | 1:1000 |
| Rodent Total Oxphos Cocktail | Mouse | Abcam, ab110413 | WB | 1:1000 |
| Sim2 | Rabbit | Abcam, ab131161 | WB, PLA | 1:500, 1:100 |
| GAPDH | Mouse | Proteintech, 60004-1-Ig | WB | 1:1000 |
| Actin | Mouse | Cell Signaling Technology, 3700S | WB | 1:1000 |
| UQCRFS1 | Rabbit | Abcam, ab191078 | PLA | 1:100 |
| anti-Chicken IgY- Alexa Fluor 488 | Donkey | Invitrogen, A78948 | IF | 1:1000 |
| anti-Goat IgG- Alexa Fluor 647 | Donkey | Invitrogen, A32849 | IF | 1:1000 |
| anti-Rabbit IgG- Alexa Fluor 568 | Donkey | Invitrogen, A10042 | IF | 1:1000 |
| anti-Guinea Pig IgG- Alexa Fluor 647 | Goat | Invitrogen, A-21450 | IF | 1:1000 |
| Anti-mouse HRP |  | Cell Signaling Technology, 7076 | WB | 1:5000 |
| Anti-rabbit HRP |  | Cell Signaling Technology, 7074 | WB | 1:5000 |

Figure S1

**G**

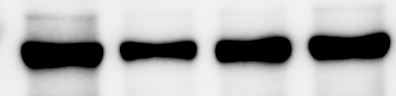

Tom70

8hD

24hD

48hD

HC11 Wild Type

Und

**A**

Und

24hrD

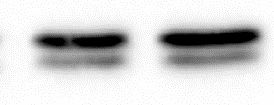

VDAC1

HC11 Wild Type

**E**

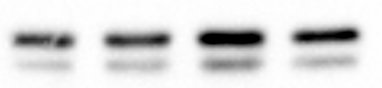

8hD

24hD

48hD

Und

VDAC1

HC11 Wild Type

**D**

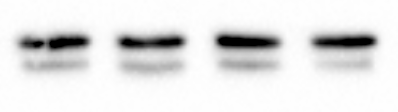

Und	8hD	24hD	48hD

VDAC1

HC11 Wild Type

**C**

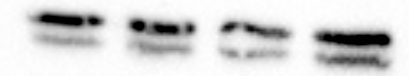

VDAC1

8hD

24hD

48hD

Und

HC11 Wild Type

Und	8hD	24hD	48hD

**B**

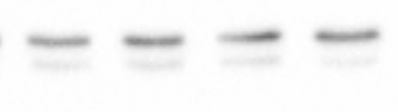

VDAC1

HC11 Wild Type

**F**

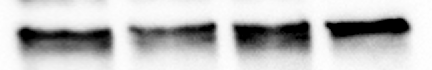

GAPDH 37kDa

Und	8hD	24hD	48hD

HC11 Wild Type

Figure S1. Even loading was ensured via parallel western blots immunoblotting for TOM70 or VDAC for mitochondrial samples. (A) Loading control for Wild Type Total OXPHOS Cocktail BN-PAGE (Fig. 2C). (B-C) Loading control for CI In Gel Activity Assay (Fig. 2E) (B) and BN-PAGE (Fig. 2G) (C). (D-E) Loading control for CIV In Gel Activity Assay (Fig. 2I) (D) and BN-PAGE (Fig. 2K) (E). (F) Loading control for CIII BN-PAGE (Fig. 2M). (G) Loading control for denaturing Total OXPHOS cocktail western using GAPDH.

Figure S2

**B**

NT Control	shParkin

Und	24hD	Und	24hD

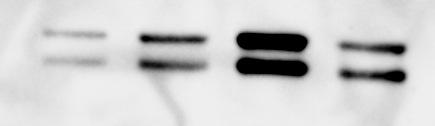

VDAC

**A**

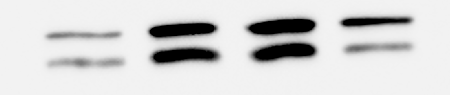

NT Control	shParkin

Und	24hD	Und	24hD

VDAC1

**C**

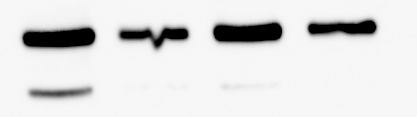

Tom70

NT Control	shParkin

Und	24hD	Und	24hD

Figure S2. Even loading was ensured via parallel western blots immunoblotting for TOM70 or VDAC for mitochondrial samples. (A) Loading control for Total OXPHOS Cocktail BN-PAGE (Fig. 3B). (B) Loading control for CI In Gel Activity Assay (Fig. 3D) (C) Loading control for CI (Fig. 3F) and CIII (Fig. 3H) BN-PAGES. CI and CIII BN-PAGES were run on the same gel using the same samples prior to immunoblotting.

Figure S3

**A**

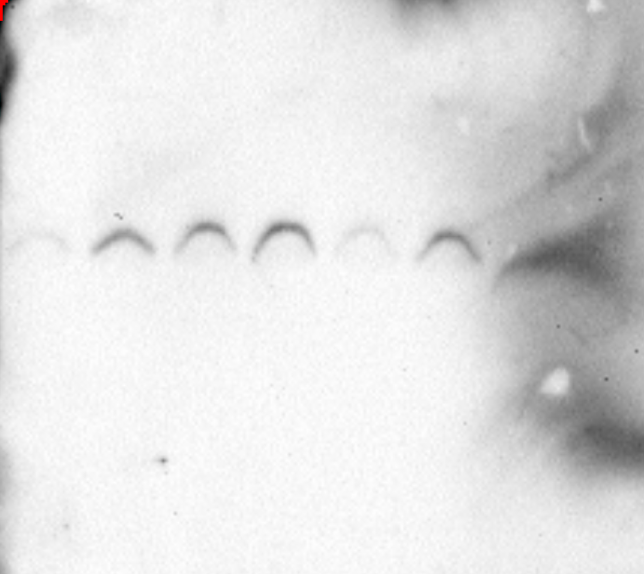

1236 kDa

1048 kDa

720 kDa

480 kDa

242 kDa

146 kDa

Scr

U

KD

U

Scr

8hD

KD

8hD

Scr

24hD

KD

24hD

Scr

48hD

KD

48hD

**shScr vs shSIM2 (KD)**

WB: SIM2

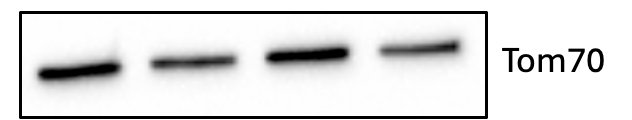

**B**

Und	8hD	24hD	48hD

HC11 Wild Type

GAPDH 37kDa

Und	8hD	24hD	48hD

Und	8hD	24hD	48hD

HC11 shScr

HC11 shSim2

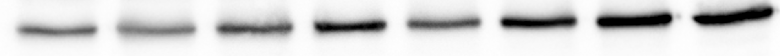

**H**

**G**

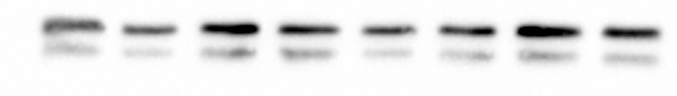

VDAC1

Und	8hD	24hD	48hD

Und	8hD	24hD	48hD

HC11 shScr

HC11 shSim2

**F**

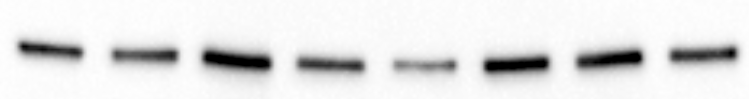

Tom70

Und	8hD	24hD	48hD

Und	8hD	24hD	48hD

HC11 shScr

HC11 shSim2

**E**

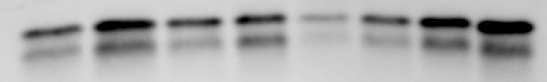

VDAC1

Und	8hD	24hD	48hD

Und	8hD	24hD	48hD

HC11 shScr

HC11 shSim2

**D**

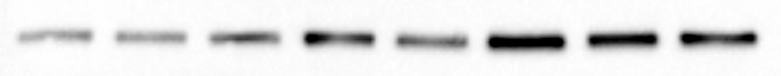

Und	8hD	24hD	48hD

Und	8hD	24hD	48hD

HC11 shScr

HC11 shSim2

Tom70

**C**

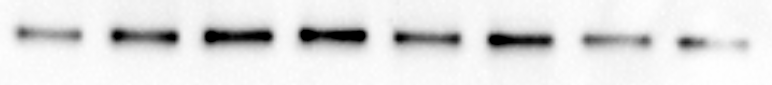

Tom70

Und	8hD	24hD	48hD

Und	8hD	24hD	48hD

HC11 shScr

HC11 shSim2

Figure S3. (A) BN-PAGE of shScr (Scr) and shSim2 (KD) differentiation time courses immunoblotting for SIM2. (B) Loading control for Wild Type SIM2 BN-PAGE (Fig. 6B). (C-D) Loading control for CI In Gel Activity Assay (Fig. 6C) (C) and BN-PAGE (Fig. 6E) (D). (E-F) Loading control for CIV In Gel Activity Assay (Fig. 6G) (E) and BN-PAGE (Fig. 6I) (F). (G) Loading control for CIII BN-PAGE (Fig. 2K). (H) Loading control for denaturing Total OXPHOS cocktail western using GAPDH.

Figure S4

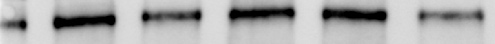

Tom70

**B**

L3

*Mito-QC*

P18

L10

L3

*Mito-QCxSim2^fl/fl^*

P18

L10

**E**

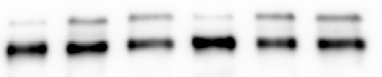

Tom70

L3

*Mito-QC*

P18

L10

L3

*Mito-QCxSim2^fl/fl^*

P18

L10

**D**

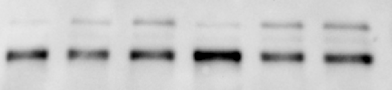

Tom70

L3

*Mito-QC*

P18

L10

L3

*Mito-QCxSim2^fl/fl^*

P18

L10

**A**

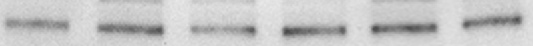

Tom70

L3

*Mito-QC*

P18

L10

L3

*Mito-QCxSim2^fl/fl^*

P18

L10

**C**

L3

*Mito-QC*

P18

L10

L3

*Mito-QCxSim2^fl/fl^*

P18

L10

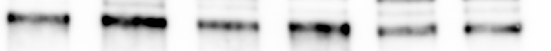

Tom70

Figure S4. Even loading was ensured via parallel western blots immunoblotting for TOM70 or VDAC for mitochondrial samples. (A-B) Loading control for CI In Gel Activity Assay (Fig. 7A) (A) and BN-PAGE (Fig. 7C) (B). (C-D) Loading control for CIV In Gel Activity Assay (Fig. 7E) (C) and BN-PAGE (Fig. 7G) (D). (E) Loading control for CIII BN-PAGE (Fig. 7I).

Figure S5

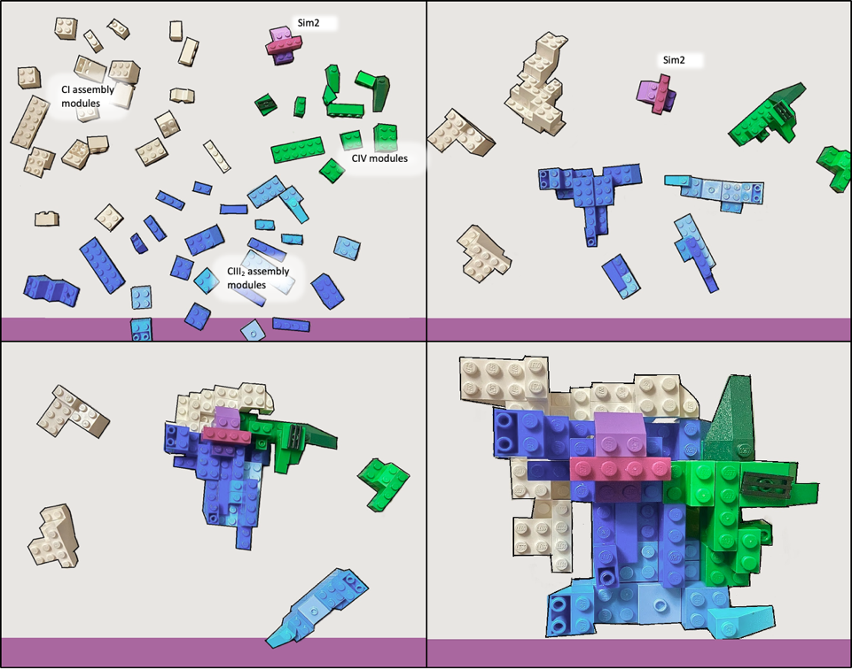

Figure S5. “Lego Model” of SC assembly demonstrating the role of SIM2 in maintaining cooperative assembly of MRC complexes.
